# Leaves in Transition: Single nuclei RNA sequencing provides insights into sorghum’s juvenile and adult phases

**DOI:** 10.64898/2026.04.22.720251

**Authors:** Oluwatoyosi M. Adaramodu, Jean G. Rosario, Erik C. Nordgren, Michel Paul, Susheel S. Bhat, Hieu Nguyen, Arif M. Ashraf, Junhyong Kim, Brent Helliker, Brian D. Gregory

**Author notes:** The author responsible for distribution of materials integral to the findings presented in this article in accordance with the policy described in the Instructions for Authors (www.plantcell.org) is Brian D. Gregory.

## Abstract

The transition from juvenile to adult phase (JA) is a key developmental process in plants, driven by conserved pathways that shape growth and stress responses. In monocots such as sorghum, this transition influences traits linked to environmental resilience, yet the regulation of micro-anatomical features such as trichomes and bulliform cells remains poorly understood. Here, we use single-nucleus RNA sequencing (snRNA-seq) to generate high-resolution gene expression maps of juvenile and adult sorghum leaves. Contrary to the traditional developmental model, we find that trichomes are present in both juvenile and adult leaves, with stage-dependent differences in gene expression. We also observe that bulliform cells, typically considered adult-specific, are present in juvenile leaves. At the transcriptomic level, cell populations cluster more strongly by developmental stage than by cell type, with juvenile cells grouping with other juvenile cells and adult cells with adult counterparts, indicating a dominant effect of developmental state on gene expression. In addition, we identify enrichment of dhurrin biosynthetic genes in trichome-associated cells as well as elevated expression of aquaporin genes in these cells, pointing to a potential role for trichomes in coordinating defense and water-related processes. Together, these findings refine current models of sorghum development and suggest that key epidermal cells and their associated functions may be established earlier than previously known. Collectively, this study provides a single nucleus resolved framework for understanding how developmental stage shapes gene expression in sorghum leaves.

## INTRODUCTION

The juvenile-to-adult (JA) phase transition is a critical developmental process in plants, marking a shift from juvenile growth to mature development (Hashimoto et al., 2019). This transition is accompanied by significant morphological, physiological, and molecular changes, enabling plants to adapt to their evolving developmental and environmental needs (Asai et al., 2002; Lawson & Poethig, 1995). In dicotyledonous plants such as *Arabidopsis thaliana* (hereafter Arabidopsis) and soybean, the JA phase transition is associated with changes in leaf morphology, trichome distribution, and shoot apical meristem (SAM) development (Telfer et al., 1997; Yoshikawa et al., 2013). These phase-specific traits are tightly regulated by conserved molecular pathways, such as those regulated by microRNA156 and microRNA172 (miR156 and miR172), which govern phase-dependent traits across a wide range of plant species (Hashimoto et al., 2019; Orkwiszewski & Poethig, 2000). Beyond morphology, juvenile-to-adult transitions are also genetically programmed and environmentally influenced (Rankenberg et al., 2021). For instance, phase-related resistance to pests and pathogens increases as the plant develops and this resistance can be seen at both whole-plant and organ levels (Develey-Rivière & Galiana, 2007; Hu & Yang, 2019). However, phase-dependent abiotic stress resilience, though equally prevalent, remains less studied despite its significance in plant adaptation.

In monocotyledonous plants such as rice, maize, and sorghum, the JA phase transition also plays a crucial role in shaping traits essential for resilience and productivity. For instance, rice transitions involve morphological changes such as slender adult leaves with pronounced midribs and a dome-shaped shoot apical meristem (SAM) (Asai et al., 2002). In maize, juvenile leaves lack bulliform cells, trichomes and display abundant epicuticular wax. By contrast, adult leaves exhibit reduced epicuticular wax and increased trichome density, particularly at the base of the leaf blade and have bulliform cells (Lawson & Poethig, 1995; Poethig, 1990; Strable et al., 2008; Bongard-Pierce et al., 1996; Orkwiszewski & Poethig, 2000). These transitions reflect critical adaptations to environmental conditions, with cells such as trichomes and bulliform cells serving distinct functional roles. For instance, bulliform cells, large vacuolated epidermal cells, facilitate leaf rolling in response to dehydration, minimizing water loss and enhancing drought resilience (Cal et al., 2019; Matschi et al., 2020). These specialized traits emphasize the importance of developmental transitions in managing abiotic stress and ensuring plant survival under fluctuating environmental conditions. However, the precise mechanisms governing the development and distribution of trichomes and bulliform cells across each developmental phase remain poorly understood in crop plants, such as sorghum.

Sorghum (*Sorghum bicolor*), a C4 plant in the Poaceae family, is particularly renowned for its exceptional drought resilience, thriving under arid and semi-arid conditions with erratic rainfall (Mundia et al., 2019). This adaptability is partly attributed to specialized traits such as bulliform cells (Al-Salman et al., 2023), which are prominent in grass leaves and facilitate leaf rolling during water stress, thereby minimizing water loss and enhancing drought tolerance (Cal et al., 2019; Matschi et al., 2020).The juvenile-to-adult (JA) phase transition in sorghum, regulated by miR156 and miR172, also influences developmental changes that shape leaf morphology and physiology (Hashimoto et al., 2019). Like maize, juvenile sorghum leaves are thought to be characterized by the absence of trichomes, with their appearance being a characteristic of adult leaves (Hashimoto et al., 2019). Relatedly, bulliform cells are well-studied in adult sorghum leaves for their role in drought adaptation. However, their developmental roles during earlier stages, particularly in juvenile leaves, remain underexplored, leaving significant gaps in understanding their contribution to sorghum’s adaptability across developmental stages.

Recent advances in single cell and single nuclei RNA sequencing (scRNA-seq and snRNA-seq, respectively), including robotics, microfluidics, and hydrogel droplets (T.-Q. Zhang et al., 2019), offer unprecedented opportunities to address these gaps. In plants, scRNA-seq has been employed to characterize cell types across various tissues in model species such as Arabidopsis, including roots, lateral roots, female gametophytes, shoot apices, leaf phloem, leaves, and stomata (Denyer et al., 2019; Illouz-Eliaz et al., 2025; Jean-Baptiste et al., 2019; Kim et al., 2021; Lee et al., 2023; Liu et al., 2020; Ryu et al., 2019; Serrano-Ron et al., 2021; Shulse et al., 2019; Song et al., 2020; Turco et al., 2019; T.-Q. Zhang et al., 2019, 2021; Hill et al., 2026; Swift et al., 2026; Tenorio Berrío et al., 2025). Despite these advancements, studies applying sc- or snRNA-seq to sorghum leaves remain limited, with most focusing on juvenile leaves and neglecting adult stages of development (Mendieta et al., 2024; Stata et al., 2025; Swift et al., 2024).

Here, we report gene expression profiling of 16,663 nuclei of sorghum juvenile and adult leaves using snRNA-seq. This approach enabled the successful isolation of gene expression profiles from nuclei of these cells, overcoming the challenges of isolating intact cells from older plant leaves for scRNA-seq (Grones et al., 2024). By identifying highly specific marker genes for each developmental phase, this method allowed us to resolve cellular heterogeneity and provided detailed insights into specialized cell types, such as trichomes, which were present in low abundances but play critical roles in the JA phase transition. We also found that trichomes and bulliform cells, traditionally associated with adult sorghum leaves, are also present in juvenile leaves, suggesting a previously unrecognized developmental continuity. Together, our results indicate that these cell types are maintained across developmental stages and may contribute to leaf structural and developmental organization in sorghum.

## MATERIALS AND METHODS

### Plant materials and growth condition

All experiments were conducted using sorghum (*Sorghum bicolor* (L.) Moench) line PI-533871, which was selected as the primary experimental line for its consistent growth and development under controlled greenhouse conditions and its widespread cultivation in the study country. Seeds were germinated in a germination mix to ensure uniform germination and then transplanted into 1/5000 Wagner pots filled with standard potting soil. Plants were grown in a controlled greenhouse environment under natural daylight, with temperature maintained at 25 ± 3°C, relative humidity at 60-70%, and a 12-hour photoperiod. Environmental conditions were monitored throughout the experiment to ensure stability. To provide comparative context and assess whether observed traits were accession specific, an additional *S. bicolor* line (PI-656076) and a wild accession (*Sorghum verticilliflorum*) were grown under the same conditions.

### Nuclei extraction

Nuclei were isolated from sorghum leaves using a protocol adapted from Guillotin et al. (2023) and Guillotin & D. Birnbaum, (2025), with slight modifications to optimize the quantity and quality of nuclei. Freshly harvested leaf sections were subjected to a predigestion step in an enzymatic solution to break down the cell walls and enhance nuclei release. The enzyme solution contained 20 mM MES, 11% D-mannitol, 3% cellulase RS, 2% macerozyme R10, 0.4% pectolyase Y-23, 2% hemicellulase, 10% viscozyme, 20 mM CaCl₂, 20 mM KCl and 0.1% BSA. The pH was adjusted to 5.8 with Tris-HCl. The leaf sections were incubated in this solution at room temperature for 10 minutes with gentle shaking.

After the digestion step, the leaf sample was washed three times in a wash buffer and immediately transferred to a pre-chilled lysis buffer to isolate the cell nuclei. The lysis buffer contained 0.3 M sucrose, 15 mM Tris-HCl (pH 8), 60 mM KCl, 15 mM NaCl, 2 mM EDTA, 0.5 mM spermine, 0.5 mM spermidine, 15 mM MES, 0.1% Triton X-100, 5 mM DTT*, 1 mM PMSF*, 1% plant protease inhibitors* (Sigma P9599), 0.4% BSA* and 0.2 µg/µL RNase inhibitor* (*indicates that the components were added immediately prior to use).

The softened tissue was finely chopped on ice with single-edged razor blades for approximately 10 minutes. The homogenized material was transferred to a pre-chilled Eppendorf tube and crushed 10 times with a pre-chilled plastic pestle. The sample was then incubated on ice for 3–5 minutes before an additional 10 strokes to ensure adequate nuclei release. The homogenate was filtered through a 20 µm mesh filter (CellTrics, 04-0042-2315) into a pre-chilled tube, with an additional 200 µL of lysis buffer used to rinse the filter.

The filtered nuclei suspension was centrifuged at 500 × g for 10 minutes at 4°C to pellet the nuclei. The pellet was resuspended in 300 µl of wash buffer (similar composition to the lysis buffer but without Triton X-100) to remove any remaining debris. After a second centrifugation step at 500 × g for 5 min, the nuclei were resuspended in 100-200 µL of a final buffer consisting of 0.3 M sucrose, 15 mM Tris-HCl (pH 8), 60 mM KCl, 15 mM NaCl, 0.5 mM spermine, 0.5 mM spermidine, 15 mM MES, 5 mM DTT*, 1% plant protease inhibitors* (Sigma P9599), 0.4% BSA* and 0.2 µg/µL RNase inhibitor*. The nuclei suspension was filtered through a 10 µm mesh filter (PluriSelect, 43-10010-50) into a clean tube to ensure purity.

To assess the quality of the isolated nuclei, an aliquot was stained with DAPI and examined under a fluorescence microscope. The concentration of nuclei was determined using a hemocytometer and the samples were adjusted to a concentration of 2,000 nuclei/µL. The freshly isolated nuclei were immediately processed for single nuclei RNA sequencing (snRNA-seq) to maintain RNA integrity and data quality.

### Single nuclei RNA sequencing (snRNA-seq) and analysis

snRNA-seq data were generated from sorghum leaves at juvenile and adult developmental stages. Juvenile and adult samples corresponded to fully expanded leaf 2 and leaf 9, respectively, selected to represent distinct heteroblastic phase identities. For each stage, nuclei were isolated from four individual plants, pooled, and partitioned into three technical channels on a Chromium Single Cell Chip (10x Genomics).

Libraries were prepared using the Chromium Single Cell 3′ Library Kit according to the manufacturer’s instructions and sequenced on a NovaSeq 6000 platform. Across the datasets, we recovered an average of ∼2,689 nuclei per sample (range: 2,009-3,042), with a mean sequencing depth of ∼201,226 reads per nucleus (range: ∼157,000-263,000). Median UMIs per nucleus were ∼609 and median genes per nucleus ∼400, with ∼18,367 genes detected per library. Raw sequencing data were processed using Cell Ranger (v7.0, 10x Genomics) to generate gene-cell count matrices, with reads aligned to the *Sorghum bicolor* reference genome (v5.1; Phytozome v13).

Gene expression matrices were analyzed using Seurat (v5.1.0). Quality control filtering was applied to retain high-quality nuclei, including thresholds of 200-5,000 detected genes and > 400 unique molecular identifiers (UMIs) per nucleus. Percentages of mitochondrial and chloroplast transcripts were calculated to assess data quality and potential organellar contamination. Given the nuclear origin of RNA in snRNA-seq, organellar transcript abundance was consistently low across samples. Mitochondrial transcript content was predominantly below 3%, and chloroplast transcript levels were negligible. As a result, no stringent filtering thresholds based on organellar transcript percentages were applied; instead, these metrics were used to confirm overall data quality and identify potential outliers.

To limit the inclusion of multiplets, nuclei with unusually low or high gene complexity were excluded based on gene and UMI thresholds. Data were log-normalized using the NormalizeData function. Highly variable genes were identified independently for each dataset using the variance-stabilizing transformation (VST) method implemented in FindVariableFeatures (nfeatures = 2000). Shared integration features across datasets were then selected using SelectIntegrationFeatures. Prior to integration, each dataset was scaled and subjected to principal component analysis (PCA) using these features. Integration anchors were identified using the FindIntegrationAnchors function with reciprocal PCA (RPCA) (reduction = “rpca”, dims = 1:30), and an integrated expression matrix containing nuclei from all samples was generated using the IntegrateData function. The integrated data were scaled and subjected to PCA, and the first 25 principal components were used to construct a shared nearest neighbor (SNN) graph using the FindNeighbors function. Clusters were identified using the FindClusters function with the Louvain algorithm at a resolution of 0.3, and UMAP was used for dimensionality reduction and visualization.

Clusters were annotated based on established marker genes and assigned to putative cell types, including mesophyll, epidermal, and trichome cells. Nuclei were annotated by developmental stage (juvenile or adult) based on sample origin. Marker gene expression was visualized using dot plots, and clustering structure was examined using UMAP projections. Cluster-enriched marker genes were identified using the FindAllMarkers function in Seurat.

Gene ontology (GO) enrichment analysis was performed using ShinyGO (v0.81) with a false discovery rate (FDR) cutoff of 0.05.

### Leaf anatomy and imaging

To examine anatomical traits, samples were collected from the central region of leaves at both juvenile and adult stages. Freshly harvested leaf tissues, excluding the mid-vein, were excised using a razor blade and fixed in either 2.5% glutaraldehyde/4% formaldehyde (EM grade) or formalin-acetic acid solution. Samples were either post-fixed in buffered osmium tetroxide or processed without post-fixation, followed by washing in distilled, deionized water. Tissues were then dehydrated through a graded ethanol series and subjected to critical point drying (Tousimis 815B, Rockville, MD, USA). Dried samples were mounted on standard 12.5 mm aluminum SEM stubs using carbon adhesive tape, with complementary adaxial and abaxial surfaces oriented upwards. Samples were sputter-coated with a 12 nm layer of platinum/palladium (60/40; Leica ACE600) and imaged using a Hitachi SU8700 scanning electron microscope (Tokyo, Japan) with appropriate detectors.

For quantitative analyses, trichome density was assessed using the nail polish impression technique (S. Wu & Zhao, 2017). Bulliform cell traits were quantified from vibratome sections. For this, leaf mid-sections (∼1cm) were excised and fixed in FPGA solution (formalin: propionic acid: glycerol: 95% ethanol: distilled water; 1:1:3:7:8). Samples were vacuum infiltrated and stored at 4 °C. Prior to sectioning, tissues were rinsed and incubated in 1× PBS for 10 min. Sections (100 µm thickness) were generated using a vibrating microtome (VT1000 S, Leica, Germany). Images were captured using a light microscope equipped with a Nikon D3300 digital camera (Japan).

### RNA isolation and quantitative RT-PCR

Total RNA was extracted from sorghum leaves using the Zymo Research RNA isolation kit, following the manufacturer’s instructions with minor modifications, including on-column DNase I treatment to remove genomic DNA contamination. RNA concentration and purity were assessed with a spectrophotometer, and integrity was confirmed by agarose gel electrophoresis. First-strand cDNA synthesis was performed using DNase-treated RNA and SuperScript III reverse transcriptase (Invitrogen).

Quantitative real-time PCR (qRT-PCR) for aquaporin genes was performed using gene-specific primers as described in Reddy et al. (2015). Each reaction was run in technical triplicates, and melting curve analysis confirmed amplification specificity. Expression levels were normalized to *ACTIN*, a commonly used internal control in plant species (Chandna et al., 2012; Gimeno et al., 2014). Three independent biological replicates were used, and relative expression values were calculated by averaging across replicates.

Quantitative RT-PCR analysis of miR156 and miR172 was performed following the method described by Hashimoto et al. (2019), using the corresponding primer sets, to validate developmental stage between juvenile and adult leaves. Eight biological replicates and three technical replicates were used for miRNA analysis. All gene-specific primers used in this study are listed in Supplementary Table S1.

### RNAscope *in situ* hybridization

RNAscope *in situ* hybridization was performed using the RNAscope 2.5 HD Duplex Detection Kit (Cat. No. 322430) according to the manufacturer’s instructions, with minor modifications. Formalin-fixed, paraffin-embedded (FFPE) sorghum juvenile leaf sections were cut at 5 µm thickness. Slides underwent target retrieval by steaming for 15 minutes. Positive and negative control probes were processed using standard manufacturer-recommended conditions. For experimental probes, amplification incubation times were adjusted for both red and green detection channels to optimize signal intensity. Slides were mounted using VectaMount mounting medium (Vector Laboratories, Cat. No. H-5000). Imaging was performed using an Aperio AT2 scanner with Z-stack focal imaging, using a Z-step interval of 0.7 µm across five focal planes.

All probe details are provided in the Supplementary Table S1.

### Data analysis

All statistical analyses and data visualizations were performed in R. For each trait, mean values and standard errors (SE) were calculated. Differences between groups were assessed using Student’s t-test. Data were visualized with bar plots, and error bars represent ± SE.

## RESULTS

### Single-nucleus transcriptome profiling identifies cell-type composition in juvenile and adult sorghum leaves

To characterize the cellular composition of sorghum leaves across developmental stages, we performed single nucleus RNA sequencing (snRNA-seq) on nuclei isolated from fully expanded leaf 2 (juvenile) and leaf 9 (adult) **(Figure 1A)**. These stages were selected to explicitly capture heteroblastic phase identity, a developmental program established early but maintained throughout leaf maturation (Bongard-Pierce et al., 1996; Poethig & Fouracre, 2024). To confirm that these leaves correspond to distinct developmental phases, we examined the expression of known phase-associated microRNAs. Consistent with previous reports (Hashimoto et al., 2019), miR156 was more highly abundant in juvenile leaves, whereas miR172 was enriched in adult leaves **(Figure 1B)**, supporting the classification of these samples as juvenile and adult. From our snRNA-seq experiment, a total of 16,663 nuclei were initially recovered, of which 11,398 passed quality filtering and were retained for downstream analysis (juvenile: 6,496; adult: 4,901). The distribution of cell types across the dataset showed that mesophyll, epidermal, and bundle sheath cells were the most abundant populations, while guard/subsidiary cells were less frequently captured **(Figure 1C)**.

**Figure 1:**
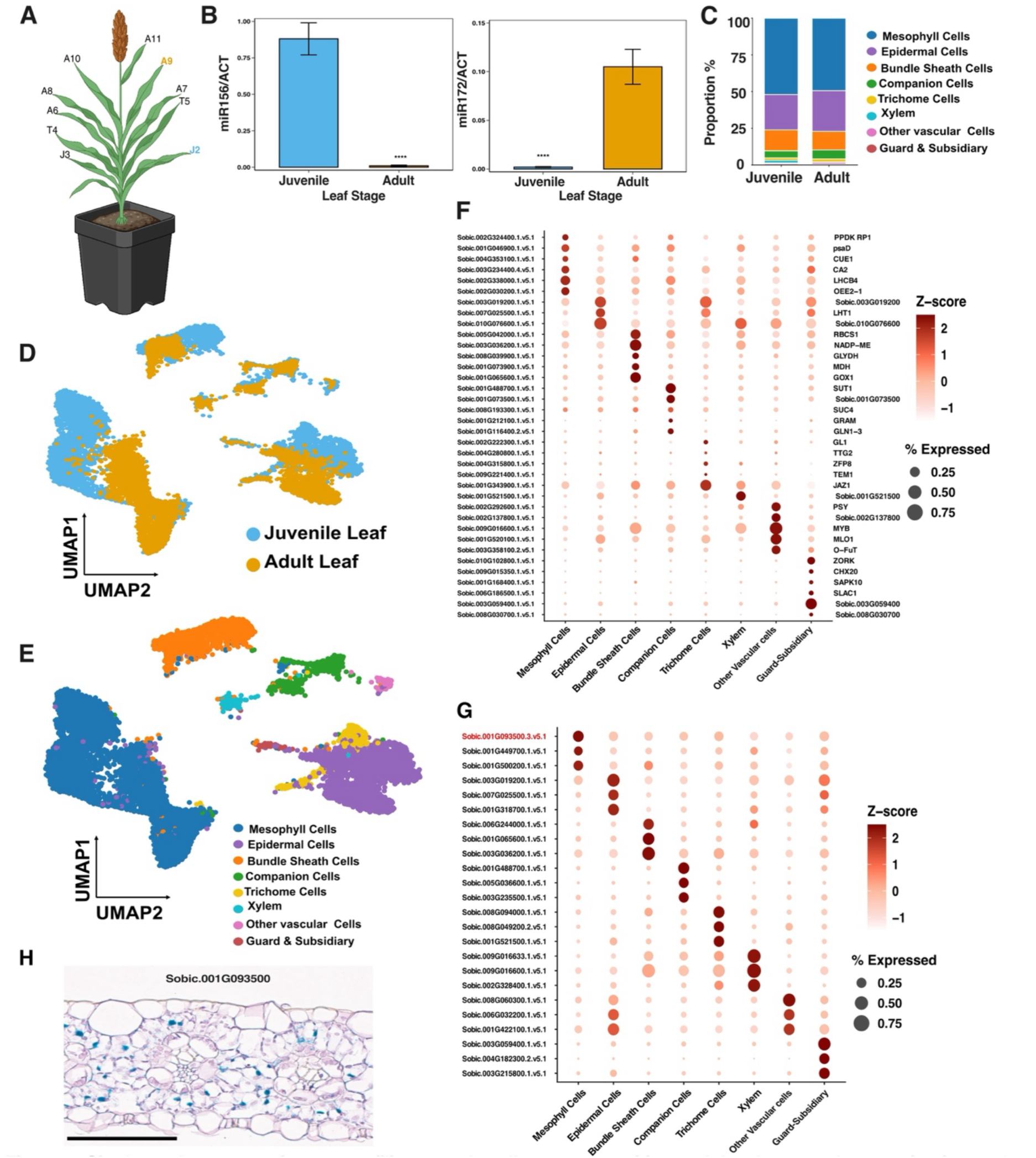
Single-nucleus transcriptome profiling reveals cell type composition and developmental separation in sorghum leaves. (A) Schematic representation of sorghum plant highlighting sampled leaves used for single nucleus RNA sequencing: fully expanded leaf 2 ouvenile, light blue (J2)) and leaf 9 (adult, orange (A9)). (B) Expression of phase-associated microRNAs showing enrichment of miR156 in juvenile leaves and miR172 in adult leaves, confirming developmental stage identity. Data are presented as mean ± s.e.m. Asterisks denote statistical significance (**** denotes p-value < 0.0001; two-tailed I-test). (C) Proportion of major cell types identified in juvenile and adult snRNA-seq datasets, including mesophyll, epidermal, bundle sheath, companion cells, trichomes, xylem, vascular cells, and guard/subsidiary cells. (D) UMAP visualization of nuclei colored by developmental stage, showing clear separation between juvenile and adult leaf populations. (E) UMAP visualization colored by annotated cell types, revealing major leaf cell populations across both developmental stages. (F) Dot plot showing expression of canonical marker genes used to annotate major cell types. Color scale represents scaled (Z-score) expression, and dot size indicates the percentage of cells expressing each gene. (G) Dot plot of selected de novo marker genes enriched in specific cell populations. The gene highlighted in red indicates the marker selected for RNAscope validation in panel (H). (H) RNAscope validation of a representative de novo marker gene *(Sobic.001G093500)* in mesophyll cells. Blue puncta indicate signal detected exclusively in mesophyll cells.. Scale bar, 20 *µm*.

To evaluate the impact of reference genome composition on cell type resolution, we performed parallel analyses using two alignment strategies: (i) utilizing a nuclear-only reference genome and (ii) a combined reference including mitochondrial and chloroplast genomes. This analysis revealed very low levels of reads mapping to both organellar genomes, as expected given our snRNA-seq focused analyses **(Supplemental Figure 1A)**. Interestingly, while overall data quality metrics remained comparable between the two approaches, inclusion of organellar genomes reduced the resolution of rare cell populations. Specifically, trichome-associated marker genes failed to form a distinct cluster when organellar genomes were included in our mapping schema **(Supplemental Figures 1B-C)**. In contrast, analysis using the nuclear-only reference retained a clearly defined trichome cluster with enriched expression of canonical trichome markers **(Supplemental Figure 1B)**. Given that mitochondrial and chloroplast transcript levels were low and within ranges for plant snRNA-seq (Grones et al., 2024) **(Supplemental Figure 1A)**, we proceeded with the nuclear-only reference for downstream analyses to preserve resolution of rare cell types.

Clustering of the single nuclei datasets revealed clear transcriptome structure. For instance, when visualized by developmental stage, nuclei separated into the two broad groups corresponding to juvenile and adult leave nuclei **(Figure 1D)**. Clustering further resolved multiple cell populations, including mesophyll, epidermal, bundle sheath, companion cells, trichomes, xylem, vascular cells, and guard/subsidiary cells **(Figure 1E)**. Nuclei from different technical channels were well distributed across clusters, with no evidence of batch-driven clustering **(Supplemental Figure 2)**.

To annotate the clusters, we combined cluster-specific gene expression patterns with Gene Ontology (GO) enrichment and comparison to publicly available plant transcriptome datasets. The expression patterns of cluster-enriched genes were cross-referenced with PlantDB (He et al., 2024), PlantCellMarker (Wendrich et al., 2020), LCMshinyapp (Fu et al., 2024), and PlantscRNAdb (Chen et al., 2021). Additionally, GO enrichment analysis was performed to further support functional annotation **(Supplemental Table 2)**. Marker prioritization was guided by sorghum-derived genes where available, followed by homologs from maize, rice, and *Arabidopsis*.

From these analyses, mesophyll clusters were characterized by the expression of genes associated with photosynthesis and carbon assimilation. These included *Sobic.002G324400* (PPDK regulatory protein 1), *Sobic.001G046900* (*psaD*), *Sobic.004G353100* (phosphoenolpyruvate/phosphate translocator 2) (Swift et al., 2024), *Sobic.003G234400* (*CA2*) (Mendieta et al., 2024), *Sobic.002G338000* (*LHCB4*) (Swift et al., 2024), and *Sobic.002G030200* (oxygen-evolving enhancer protein 2-1) (Swift et al., 2024). Consistent with the expression profile, GO enrichment analysis highlighted terms related to photosynthesis and photosystem II, supporting the identification of this cluster as mesophyll cells. In contrast, bundle sheath clusters showed a distinct expression pattern associated with C_4_ metabolism. For instance, genes such as *Sobic.003G036200* (*NADP-DEPENDENT MALIC ENZYME*) (Chang et al., 2012), *Sobic.001G073900* (*MALATE DEHYDROGENASE*) (Mendieta et al., 2024), *Sobic.008G039900* (GLYCINE *DEHYDROGENASE*) (Döring et al., 2016), *Sobic.001G065600* (*GLYCOLATE OXIDASE 1*) (Mendieta et al., 2024), and *Sobic.005G042000* (*RBCS1*) (Mendieta et al., 2024) were enriched in this cluster, consistent with the role of bundle sheath cells in CO₂ concentration and fixation **(Figure 1F)**.

Epidermal clusters were defined by the expression of *Sobic.007G025500* (*LYSINE HISTIDINE TRANSPORTER 1*) (Marand et al., 2021). In addition, transcripts such as *Sobic.003G019200* (ortholog of *Zm00001d008923*) and *Sobic.010G076600* (ortholog of *Zm00001d044995*; *SnRK*) were consistently enriched in this cluster, although their functions are currently unannotated (Marand et al., 2021). Companion cell clusters were readily identifiable through the expression of known sucrose transporters, including *Sobic.001G488700* (*SUT1*) (Matsukura et al., 2000; Scofield et al., 2006) and *Sobic.008G193300* (*SUC2*) (Lee et al., 2023; Truernit & Sauer, 1995), which are known to play roles in phloem loading and sugar transport. Similarly, trichome clusters exhibited expression of genes with previously characterized roles in *Arabidopsis*, including Sobic.002G222300 (*GLABROUS1*), *Sobic.004G280800* (*TRANSPARENT TESTA GLABRA2*), *Sobic.004G315800* (*ZFP8*), *Sobic.009G221400* (*TEM1*), and *Sobic.001G343900* (*JAZ1*) (Gan et al., 2007; Larkin et al., 1994; Matías-Hernández et al., 2016; Qi et al., 2011; Y. Wang et al., 2022; Z. Wang et al., 2019) **(Figure 1F)**. These genes have also been linked to trichome development and regulation. Guard and subsidiary cell populations were identified based on the shared expression of genes involved in ion transport and signaling. These included *Sobic.006G186500* (*SLAC1*) (Min et al., 2019), *Sobic.010G102800* (*SKOR*) (T. H. Nguyen et al., 2017; Stata et al., 2025), *Sobic.001G168400* (*SAPK10*) (Min et al., 2019), *Sobic.009G249900* (*GH3 DOMAIN-CONTAINING PROTEIN*) (Marand et al., 2021), and *Sobic.009G015350* (*CATION/H⁺ ANTIPORTER 20*) (Sun et al., 2022), all of which are associated with stomatal function and cellular ion balance. Finally, xylem and other vascular-associated clusters showed enrichment of genes such as *Sobic.004G062500* (*BM5*), *Sobic.002G292600* (*PHYTOENE SYNTHASE*), *Sobic.004G062500* (*4CL3*), *Sobic.009G016600* (*MYB TRANSCRIPTION FACTOR*), *Sobic.007G047300* (*FLAVANOL 3-O-METHYLTRANSFERASE*), *Sobic.001G520100* (*MLO1*), *Sobic.005G139700* (*RBOHD*), *Sobic.003G358100* (*O-FUCOSYLTRANSFERASE*), and *Sobic.005G186800* (*HYDROXYMANDELONITRILE LYASE*) (Stata et al., 2025), suggesting roles in vascular-associated metabolism and defense **(Figure 1F; Supplemental Figure 3A)**.

Importantly, the combination of literature marker gene enrichment, GO enrichment, **(Supplemental Tables 2-3)**, and comparison to existing plant datasets supported the assignment of major leaf cell types. In addition to previously described marker genes, we identified de novo markers enriched within specific clusters across cell types and development stage **(Figure 1G; Supplemental Figure 3B; Supplemental Tables 4-5)**, which provides further resolution of cell identity for future analyses of cell type-specific gene expression studies in sorghum leaves. Importantly, one of the newly identified marker genes (*Sobic.001G093500.3*) for sorghum leaf mesophyll cells was validated as specific to this cell type using RNAscope **(Figure 1H)**. In total, our results identified the major cell types of sorghum juvenile and adult leaves, including identifying new gene markers for each identify.

### Trichomes are present in both juvenile and adult sorghum leaves

Trichomes have traditionally been described as phase-change traits in grasses, associated predominantly with the adult developmental stage (Doody et al., 2022; Hashimoto et al., 2019; Poethig, 1990; Telfer et al., 1997; Y. Xu et al., 2019). In contrast to this view, our snRNA-seq analysis identified a distinct trichome cell cluster in both juvenile and adult sorghum leaves **(Figures 1C, 1E, and 2A)**. This cluster was defined by the expression of established trichome marker genes, supporting its cellular identity **(Figure 2B).** Notably, although a greater number of trichome cells was captured in juvenile leaves, normalization by total cell number revealed that trichomes constituted 1.6% of juvenile cells and 2.0% of adult cells **(Figure 2A)**. Further analysis of the top group-enriched marker genes identified using *FindAllMarkers* within the trichome subset further showed that juvenile and adult trichome cells exhibit stage-biased expression patterns including genes associated with transcription regulation (e.g., WRKY and CBF family members), transport processes (e.g., ABC transporter family proteins), and protein kinase signaling **(Figure 2C; Supplemental Table 6).** Trichome-enriched genes were identified by performing differential expression analysis within the trichome subset (juvenile vs. adult) using *FindAllMarkers*, and selecting the top-ranking genes based on log fold change. Overall, these results support the idea that trichomes are present on both juvenile and adult sorghum leaves, but the overall gene expression profiles of these cells are quite different between these different leaf stages.

**Figure 2:**
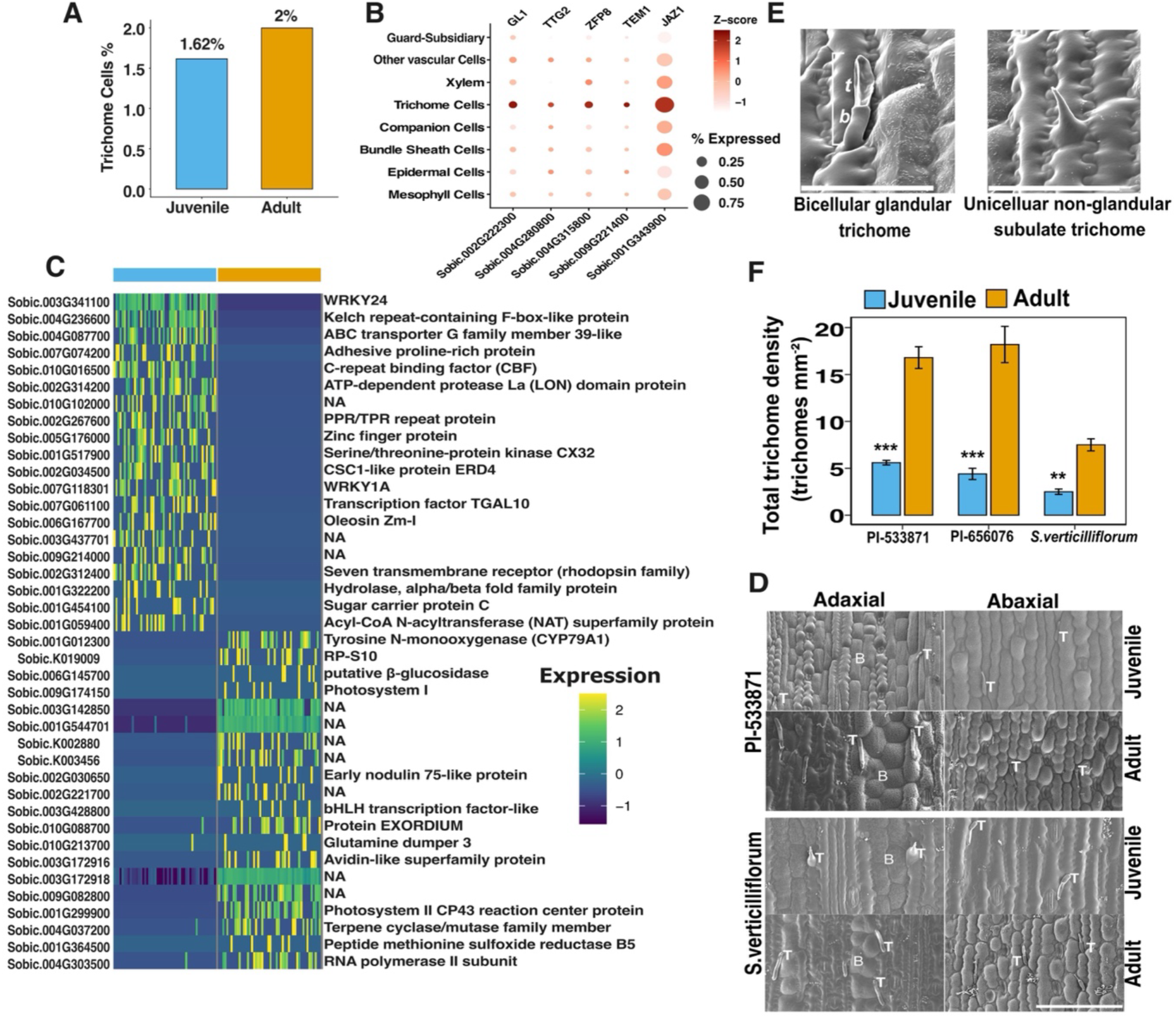
Trichomes are present in both juvenile and adult sorghum leaves. **(A)** Proportion of trichome cells identified from snRNA-seq data in juvenile and adult leaves, showing comparable representation across developmental stages after normalization by total cell number. (B) Dot plot showing expression of established trichome marker genes across major cell types. Color scale represents scaled (Z-score) expression, and dot size indicates the percentage of cells expressing each gene. (C) Heatmap of top stage-enriched genes identified within the trichome cluster, highlighting distinct transcriptional profiles between juvenile and adult trichome cells. Genes were selected based on differential expression analysis (FindAIIMarkers). “NA” indicates genes without annotated gene names in the Phytozome database. (D) Scanning electron microscopy (SEM) images of leaf epidermal surfaces showing trichomes on both adaxial and abaxial sides in juvenile and adult leaves across sorghum lines. T, trichome; B, bulliform cell. Scale bar, 100 *µm.* (E) Representative SEM images showing trichome morphology, including bicellular glandular trichomes and unicellular non-glandular trichomes. Scale bar, 100 *µm* (F) Quantification of total trichome density (combined adaxial and abaxial surfaces) across sorghum lines, showing increased density in adult leaves relative to juvenile leaves. Data are presented as mean± s.e.m. Asterisks denote statistical significance (** denotes p-value _<_ 0.01 and*** denotes p-value _<_ 0.001; two-tailed t-test) N = 10.

To validate these whole transcriptome-based observations, we examined leaf epidermal surfaces using scanning electron microscopy (SEM). This analysis revealed that trichomes were clearly observed on both adaxial and abaxial surfaces in juvenile and adult leaves **(Figure 2D).** Morphologically, these structures included bicellular glandular trichomes, consisting of a basal cell and an apical secretory cell, as well as unicellular non-glandular trichomes **(Figure 2E),** consistent with previous descriptions in other plants (Triplett et al., 2023; Watts & Kariyat, 2021). To determine whether this trichome pattern is influenced by domestication, we examined both leaf surfaces of juvenile and adult leaves of a wild sorghum parental line. Specifically, we used *Sorghum verticilliflorum* for this analysis and found that it exhibited the same qualitative pattern, with trichomes present in both juvenile and adult leaves. We further quantified total trichome density by combining counts from the adaxial and abaxial leaf surfaces **(Figure 2F)** and found that juvenile leaves consistently exhibited lower trichome density compared to adult leaves across all three ecotypes examined, with the wild line displaying lower levels of trichomes at both leaf developmental stages as compared to the two domesticated ecotypes. Taken together, these results indicate that trichomes are not restricted to the adult phase in sorghum but are instead present across developmental stages. This conserved pattern across the three lines refines our current understanding of trichome development in grasses and raises the possibility that trichomes may contribute to leaf function throughout development, including potential roles in environmental response in sorghum.

### Trichome-associated enrichment of the dhurrin biosynthetic pathway in sorghum leaves

To investigate the functional specialization of cell types in sorghum leaves, we performed Gene Ontology (GO) enrichment analysis for each cell cluster **(Supplementary Table 2)**. Notably, we found that the trichome cluster was enriched for terms related to dhurrin biosynthesis, including cyanogenic glycoside biosynthetic process and dhurrin biosynthetic process **(Figure 3A)**. Dhurrin, a cyanogenic glucoside, is abundant in *Sorghum bicolor* and represents a key biochemical feature distinguishing sorghum from other C₄ grasses such as maize, millet, and sugarcane (B. Wang et al., 2024). This observation prompted us to examine the dhurrin biosynthetic pathway in the context of cell-type-specific expression.

**Figure 3:**
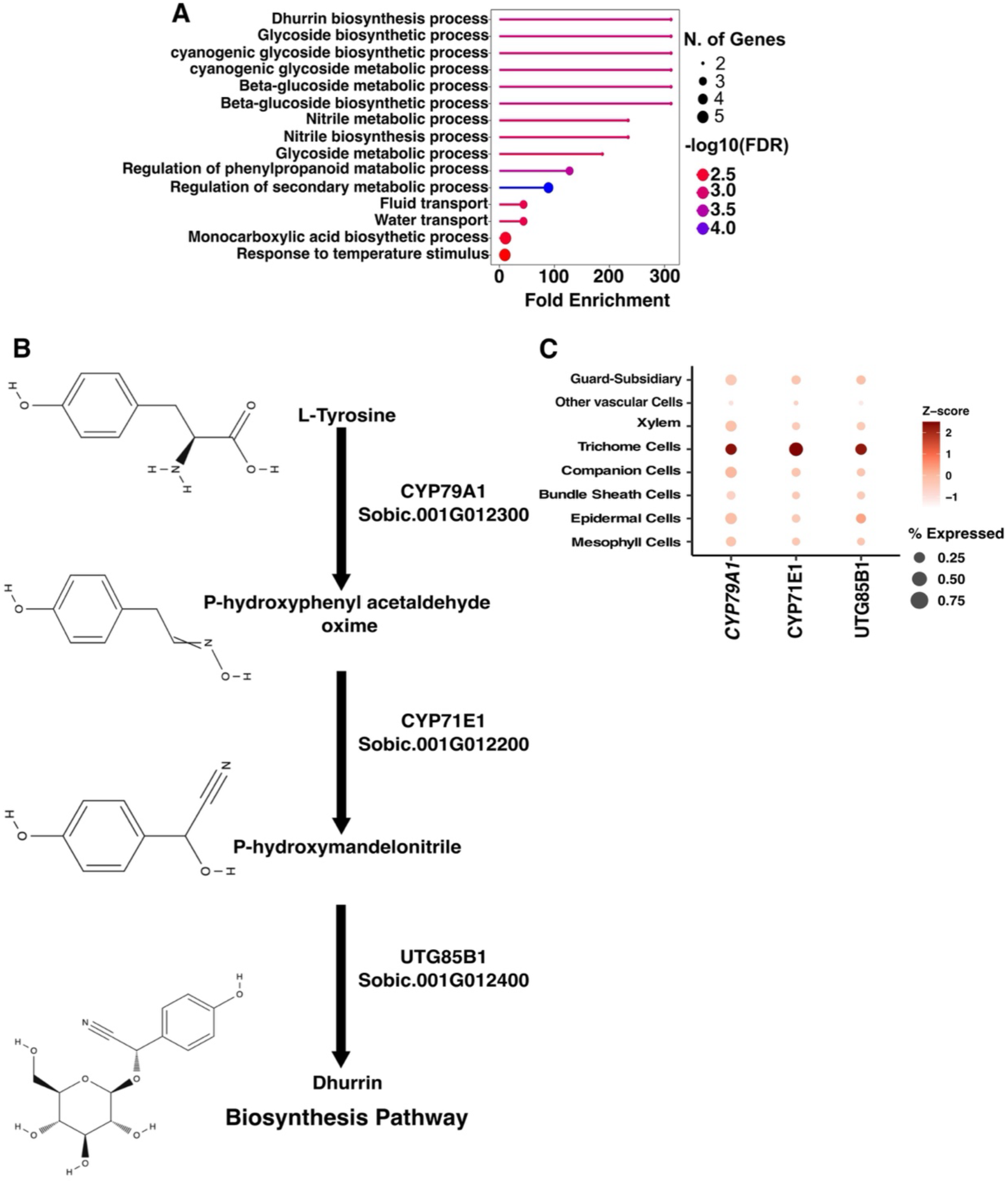
Dhurrin biosynthesis is enriched in trichome-associated cell populations. **(A)** Gene Ontology (GO) enrichment analysis of genes expressed in trichome-associated cells, highlighting overrepresentation of pathways related to cyanogenic glucoside biosynthesis, glycoside metabolism, and water transport. Dot size indicates the number of genes, and color represents significance (-log10 FDR). (B) Schematic of the dhurrin biosynthesis pathway in Sorghum bicolor, showing the stepwise conversion of L-tyrosine to p-hydroxyphenylacetaldehyde oxime, p-hydroxymandelonitrile, and finally dhurrin. Each reaction is catalyzed by a dedicated enzyme: CYP79A1 *(Sobic.001G012300),* CYP71E1 *(Sobic.001G012200),* and UGT85B1 *(Sobic.001G012400).* (C) Dot plot showing the expression of key dhurrin biosynthetic genes (CYP79A1, CYP71E1, and UGT85B1) across annotated cell types. Color scale represents scaled (Z-score) expression, and dot size indicates the percentage of cells expressing each gene. Enrichment of these genes is observed in trichome cells relative to other cell types.

Dhurrin biosynthesis proceeds from L-tyrosine through a three-step enzymatic pathway **(Figure 3B**). CYP79A1 catalyzes the conversion of L-tyrosine to p-hydroxyphenylacetaldehyde oxime, which is subsequently converted to p-hydroxymandelonitrile by CYP71E1. Finally, UGT85B1 mediates the glucosylation of this second intermediate to form dhurrin (Cowan et al., 2022; Darbani et al., 2016; Koleva et al., 2025; Wang et al., 2024). Thus, we interrogated whether the enrichment of this pathway in the trichome cell cluster was driven by the expression of a single key enzyme or whether the entire biosynthetic pathway was co-localized within this cell type. Our single-nucleus atlas showed that all three core enzymes are quite highly expressed in the trichome cell cluster **(Figure 3C)**. Specifically, we found that *CYP79A1* displayed strong expression in trichome cells, while *CYP71E1* and *UGT85B1* were also enriched in this cluster, with comparatively lower expression observed in other cell types. These results suggest that multiple steps of the dhurrin biosynthetic pathway are most significantly associated with trichome cells in sorghum leaves.

### Bulliform cells are present in both juvenile and adult leaves of sorghum plants

Bulliform cells (BCs) are specialized, vacuolated cells located on the adaxial surface of grass leaves, forming clusters along the length of the leaf. In maize, these cells are arranged in longitudinal strips of two to five cells and are exclusively found in the adaxial epidermis (Becraft et al., 2002; Matschi et al., 2020; Sylvester & Smith, 2009). Their appearance in maize is a prominent marker of vegetative phase change, signifying the transition from juvenile to adult leaves (Bongard-Pierce et al., 1996; Vega et al., 2002). Furthermore, BCs have been hypothesized to facilitate leaf rolling through changes in turgor pressure, enabling leaves to reduce surface exposure and limit water loss during drought conditions (Matschi et al., 2020; Moulia, 2000). Our finding that juvenile sorghum leaves displayed trichomes, a trait often associated with the adult phase, prompted us to reexamine the presence of bulliform cells (BCs) in juvenile leaves. To do this, we used scanning electron microscopy (SEM) analysis of leaf cross sections from domesticated (PI-533871 and PI-656076) and wild (*Sorghum verticilliflorum*) sorghum lines. This analysis revealed the presence of BCs in both juvenile and adult sorghum leaves from both domesticated and wild ecotypes **(Figures 4A-B)**. Interestingly, we also found that trichomes are frequently observed near bulliform cells on the adaxial leaf surface **(Figure 4C)**, suggesting a potential spatial association between these two cell types. This observation suggests that unlike in maize where bulliform cells are reported to be absent in juvenile leaves, bulliform cells in sorghum are present across both developmental stages.

**Figure 4:**
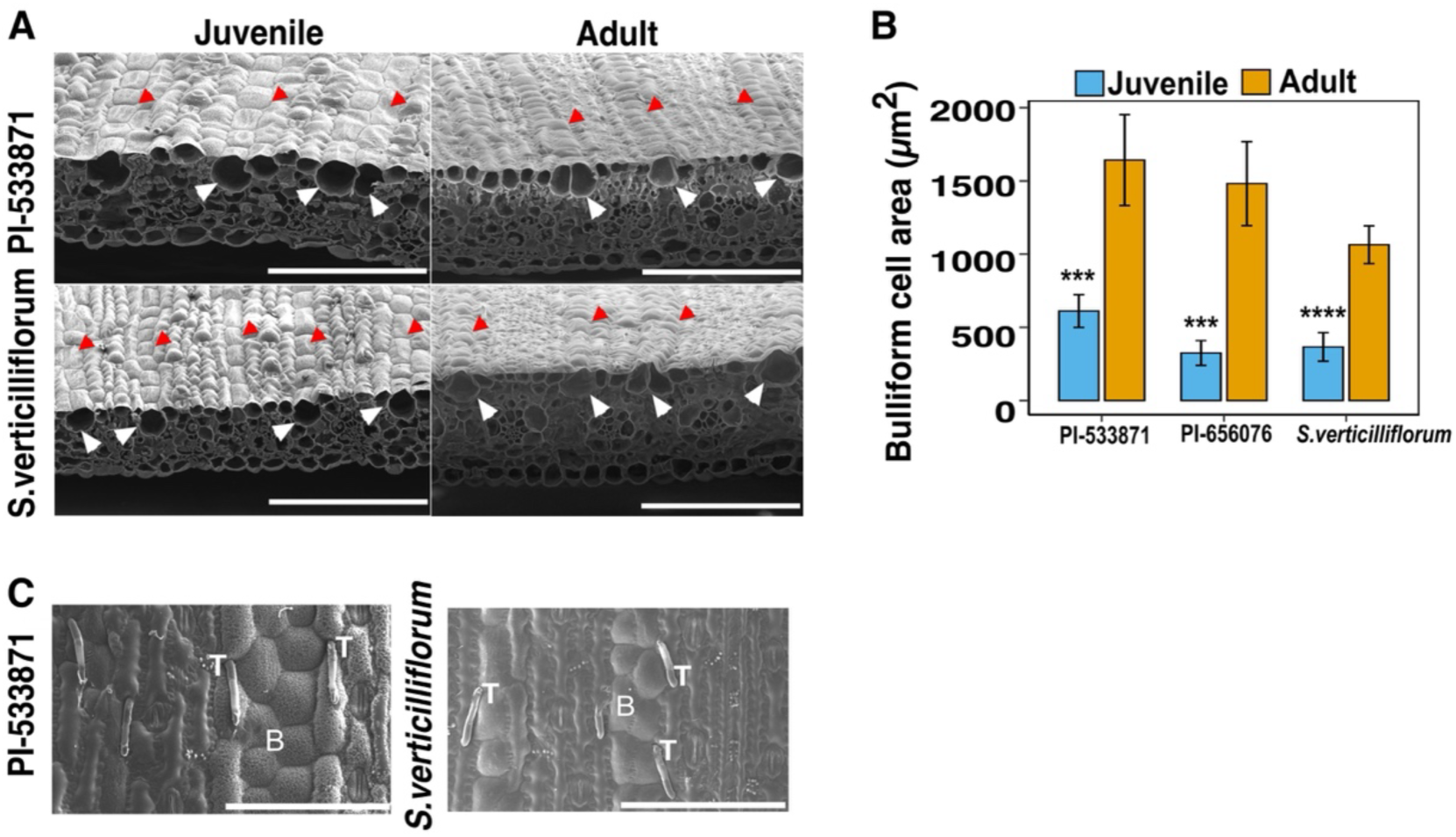
Bulliform cells are present in both juvenile and adult sorghum leaves. (A) Scanning electron microscopy (SEM) images of juvenile and adult leaves from domesticated (Pl-533871) and wild (S. verticilliflorum) sorghum lines. Red arrowheads indicate bulliform cells on the adaxial leaf surface, while white arrowheads highlight bulliform cells in cross-sectional view. Scale bars, 100 *µm.* (B) Quantification of bulliform cell area across sorghum lines, showing increased cell size in adult leaves relative to juvenile leaves. Data are presented as mean± s.e.m. Asterisks denote statistical significance (*** denotes p-value _<_ 0.001 and •••• denotes p-value _<_ 0.0001; two-tailed I-lest) **1\1=** 10. (C) SEM images of the adult adaxial leaf surface showing the spatial relationship between trichomes (T) and bulliform cells (BJ in the specified ecotypes of sorghum. Scale bars, 100 *µm*.

### Developmental stage shapes transcriptome identity in sorghum leaves

To assess the extent of shared and stage-specific transcriptome programs in sorghum leaves, we first compared genes detected in juvenile and adult leaves. Genes were classified based on their presence or absence in each developmental stage and we found that majority of genes (12,790) were expressed in both juvenile and adult leaves, indicating a largely conserved transcriptional framework **(Figure 5A)**. In contrast, 2,801 genes were uniquely detected in juvenile leaves, while 1,322 genes were specific to adult leaves. These values were derived by identifying genes with detectable transcript counts in each stage and comparing their overlap **(Figure 5A)**. These results indicate that while most genes are shared across developmental stages, each stage is also associated with a distinct subset of expressed genes, suggesting stage-dependent transcriptional specialization.

**Figure 5:**
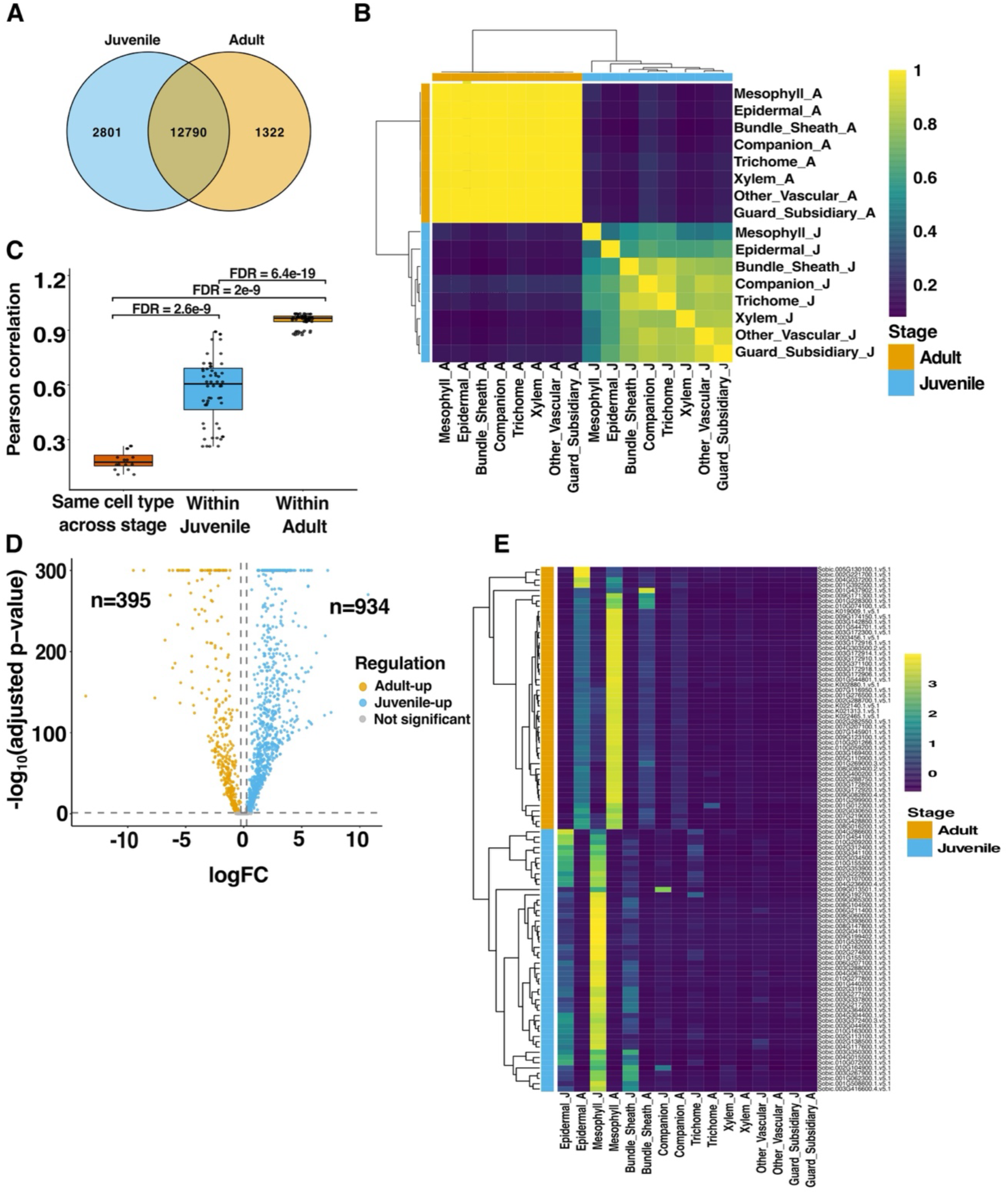
Developmental stage dominates transcriptional organization in sorghum leaves. (A) Venn diagram showing overlap of expressed genes between juvenile and adult leaves. Most genes are shared across stages, with a subset uniquely detected in either juvenile or adult samples. (B) Hierarchical clustering heatmap of transcriptome similarity across cell types and developmental stages based on averaged gene expression profiles. Samples cluster primarily by developmental stage ijuvenile versus adult) rather than by cell type. Color scale represents pairwise correlation. (C) Quantification of pairwise transcriptome similarity. Pearson correlation coefficients are shown for the same cell type across stages, and for different cell types within juvenile and adult leaves. Correlations are higher within developmental stages than across stages. (FDR values indicated). (D) Volcano plot showing differential gene expression between juvenilE and adult leaves. Genes upregulated in juvenile (blue) and adult (yellow) stages are indicated, with significantly differentially expressed genes highlighted (adjusted P value threshold). (E) Heatmap of top stage-enriched genes across all annotated cell types. Genes were selected based on differential expression analysis, revealing coordinated and stage-specific expression patterns across multiple cell populations.

We next examined how these transcriptional differences are organized across cell types. To this end, we performed hierarchical clustering of transcriptome profiles across all annotated cell populations and developmental stages. Average gene expression profiles were computed for each cell type within each stage, and pairwise similarity was assessed to generate a clustering dendrogram. Notably, samples clustered primarily by developmental stage rather than by cell type **(Figure 5B)**. For example, mesophyll, epidermal, and vascular-associated cells from juvenile leaves grouped more closely with other juvenile cell types than with their corresponding adult counterparts. This pattern suggests that developmental stage exerts a strong influence on transcriptional identity, shaping gene expression profiles across diverse cell populations. To further quantify this observation, we calculated pairwise transcriptome correlations across cell types and developmental stages. Correlation coefficients were computed using averaged expression values across genes for each cell type within each stage. Cells of the same type across different stages exhibited lower similarity compared to cells within the same developmental stage. Specifically, correlations between juvenile and adult versions of the same cell type were consistently lower than correlations among different cell types within either juvenile or adult leaves **(Figure 5C)**.

We then sought to identify the genes contributing to these stage-dependent differences. Differential expression analysis was performed using *FindMarkers* on the integrated dataset to identify genes enriched in juvenile versus adult leaves. We identified a substantial number of genes upregulated in each developmental stage **(Figure 5D)**. To determine whether these abundance differences reflect structured gene programs rather than isolated gene-level changes, we examined the expression patterns of the top 50 stage-enriched genes across all cell types. These genes were selected based on their relative enrichment and fold-change values in each developmental stage. Heatmap analysis revealed distinct and coordinated expression patterns associated with each stage **(Figure 5E).** Notably, both juvenile and adult gene sets were most strongly expressed in mesophyll and epidermal cells, with comparatively weaker but detectable expression in bundle sheath cells. This observation suggests that these cell types are the major contributors to developmental transcriptome remodeling in sorghum leaves.

### Aquaporin expression is developmentally regulated and enriched in trichome cells

Having established that developmental stage strongly shapes transcriptional identity across cell types, we next examined aquaporin (*AQP*) gene expression. As a drought-tolerant species, sorghum has been widely studied for its water-use efficiency, making aquaporins (key regulators of transmembrane water transport) a relevant focus. Aquaporins (AQPs) from the PIP, TIP, and NIP subfamilies are widely implicated in plant drought responses, where they regulate transmembrane water transport and contribute to physiological plasticity under water-limiting conditions (Afzal et al., 2016; Byrt et al., 2023; Y. Li et al., 2025; Reddy et al., 2015; F. Wu et al., 2015; Zupin et al., 2017). Based on these previous studies, we examined the expression of these selected drought-associated aquaporin genes across sorghum leaf development and cell types to assess whether their expression is developmentally regulated. To first assess developmental patterns, we examined aquaporin expression across juvenile and adult leaves. Members of the PIP, TIP, and NIP families all displayed higher expression levels in juvenile relative to adult leaves **(Figure 6A)**. This pattern was observed across multiple aquaporin subfamilies, indicating a developmental bias toward higher expression in juvenile leaves.

**Figure 6:**
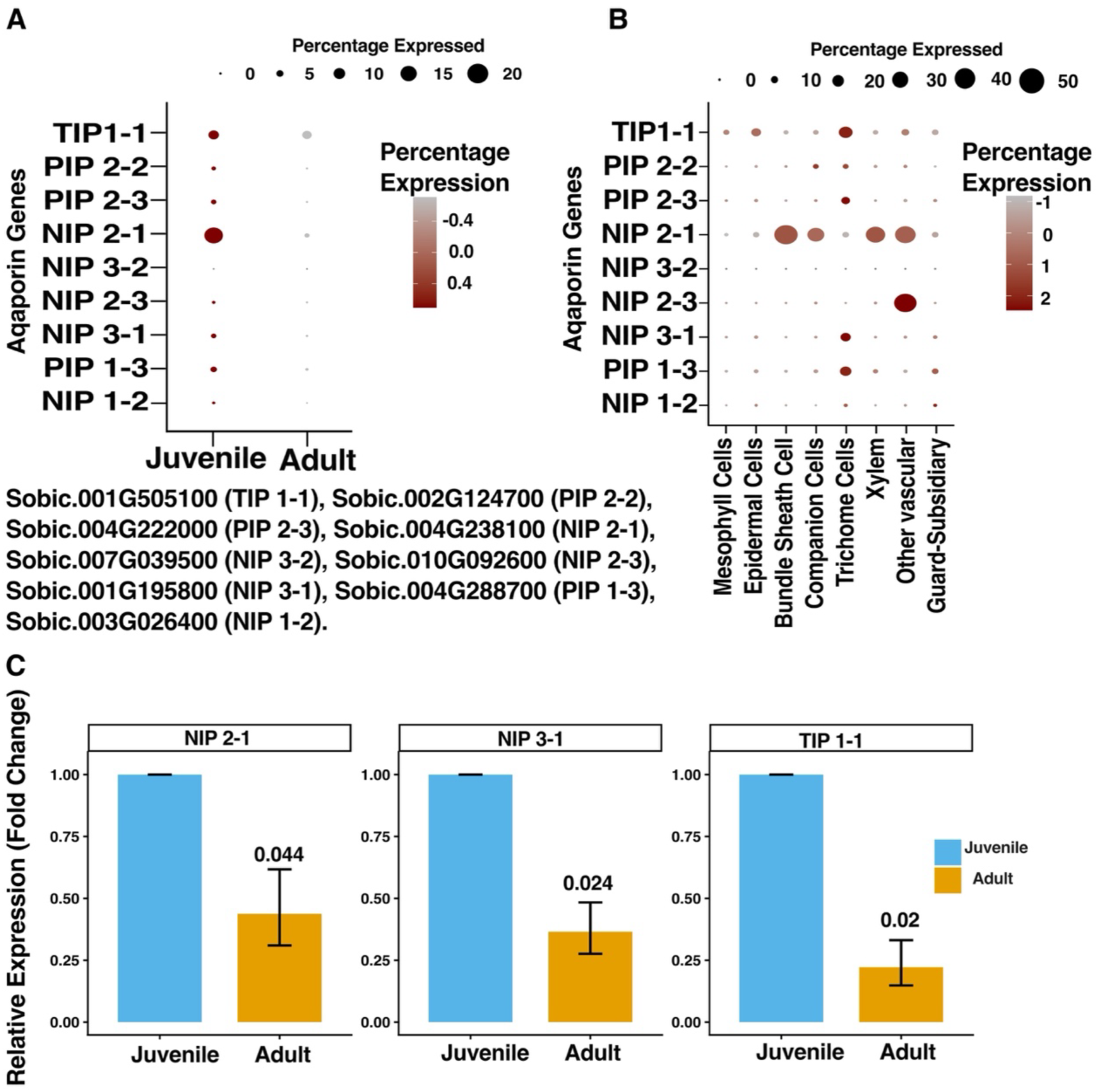
Aquaporin genes show stage-dependent expression and enrichment in specific cell types. (A) Dot plot showing expression of aquaporin (AQP) genes across developmental stages Uuvenile and adult leaves). Color scale represents average expression, and dot size indicates the percentage of cells expressing each gene. Several AQPs display relatively higher expression in juvenile leaves compared to adult leaves. (B) Dot plot showing expression of AQP genes across annotated cell types. Color scale represents average expression, and dot size indicates the percentage of cells expressing each gene. (C) Quantification of selected AQP genes (NIP2-1, NIP3-1, and TIP1-1) showing relative expression (fold change) between juvenile and adult leaves. Data are presented as mean± s.e.m. P-values are indicated above bars (Two-tailed t-test).

We next examined the distribution of aquaporin expression across cell types. This analysis revealed pronounced cell type specificity, with distinct enrichment patterns observed across the leaf **(Figure 6B)**. Notably, trichome cells showed enrichment of multiple plasma membrane intrinsic proteins (PIPs), including members of the PIP1 and PIP2 clades, as well as *TIP1-1* and NIP family members. This enrichment pattern distinguishes trichomes from other epidermal and vascular-associated cell types. In contrast, some aquaporins showed preferential enrichment in vascular-associated cell types rather than trichomes, highlighting heterogeneity in cell type–specific expression within the aquaporin family. To further validate these developmental patterns, we performed targeted expression analysis of selected genes, including *NIP2-1*, *NIP3-1*, and *TIP1-1* in juvenile as compared to adult leaves using qRT-PCR analyses. Using this analysis, we found that these genes showed significantly higher levels in juvenile leaves as compared to adult leaves as expected **(Figure 6C)**, supporting the developmental trends observed in our global snRNA-seq analysis. We were also able to validate the cell type specific expression of TIP1-1 in BCs and trichomes, which are found in the same cluster in our snRNA-seq analysis, using RNAScope analysis **(Supplemental Figure 4)**. In total, these findings reveal that *AQP* transcripts demonstrate a highly specific developmental and cell type-specific expression pattern, which might play into the significant environmental resilience displayed by sorghum plants. This hypothesis will require further testing in the future.

## DISCUSSION

The transition from juvenile to adult phases in plants governs the development of traits essential for environmental adaptation, stress resilience, and reproductive success (Bongard-Pierce et al., 1996; Hashimoto et al., 2019). In this study, we present novel insights into the molecular and anatomical changes that occur during the juvenile-to-adult phase transition in sorghum, emphasizing the roles of trichomes, bulliform cells, and aquaporins across developmental stages.

### The role of miR156 and miR172 in plant phase transition

Our study confirms that, like in other plants, sorghum exhibits similar expression patterns of miR156 and miR172, which are essential regulators of the juvenile-to-adult transition. These microRNAs are well-documented in species such as Arabidopsis and maize, where they govern key developmental changes that differentiate juvenile and adult phases (Hashimoto et al., 2019; Poethig, 2003, 2013). We found that in sorghum, as in other plants, the expression of miR156 and miR172 displays a developmental stage specific expression pattern **(Figure 1B)**, suggesting that like in other plants these miRNAs play crucial roles in influencing the shift from juvenile to adult leaf characteristics.

### Developmental persistence and metabolic specialization of trichomes in sorghum leaves

In *Arabidopsis*, trichomes are a key marker of the adult phase, emerging on the abaxial surface of leaves during the juvenile-to-adult transition. This process is regulated by transcription factors such as GLABRA1 (GL1) and GL3, which promote trichome formation once a developmental threshold is reached (Huijser & Schmid, 2011; Poethig, 2003). Similarly, in maize, trichomes are absent in juvenile leaves but appear in adult leaves, primarily at the base of the leaf blade, where they have been associated with protection against environmental stressors such as drought and herbivory (Okamoto et al., 2020; Poethig, 1990). Previous studies in sorghum have suggested that trichomes are absent from juvenile leaves, consistent with the phase transition model observed in maize and Arabidopsis (Hashimoto et al., 2019). However, our findings, based on snRNA-seq and anatomical analyses, indicate that trichomes are present in both juvenile and adult sorghum leaves, although they are more abundant in the adult stage. This pattern was consistently observed across the two domesticated sorghum lines examined, as well as in a wild relative, using both snRNA-seq and SEM **(Figure 2)**. Based on the images presented in prior studies (Hashimoto et al., 2019; Yoshikawa et al., 2013), trichome identification appears to rely on surface-level visualization approaches, which may have limited sensitivity for detecting low-density or small structures at early developmental stages. Such approaches may also be biased toward more prominent, hair-like trichomes, while shorter or less conspicuous trichomes may be more difficult to resolve. In contrast, SEM provides higher-resolution visualization of epidermal features, and in our study enabled the detection of trichomes on both adaxial and abaxial surfaces in juvenile and adult leaves **(Figure 2D)**. Together, these observations suggest that, rather than being strictly absent during early development, trichomes in sorghum may be established earlier but at lower abundance or with less conspicuous morphology, becoming more pronounced as leaves mature.

This developmental persistence raises the possibility that trichomes in sorghum are not merely markers of phase identity, but functionally relevant throughout leaf development. In this context, the enrichment of key enzymes involved in dhurrin biosynthesis within trichome cell suggests a spatial organization of cyanogenic metabolism **(Figure 3)**. Dhurrin, a cyanogenic glucoside, plays a central role in plant defense and nitrogen management in *Sorghum bicolor*, and has also been linked to physiological traits such as water-use efficiency and environmental adaptation (Morris et al., 2026). While previous studies have described dhurrin as broadly distributed across the epidermis (Thayer & Conn, 1981), our observations point to a more localized epidermal cell type expression pattern **(Figure 3)**. Such spatial restriction may provide a physiological advantage, as cyanogenic metabolism is both energetically costly and potentially auto-toxic (Bjarnholt et al., 2018). Localizing biosynthesis to trichomes **(Figure 3)** could allow sorghum to maintain an effective chemical defense barrier at the leaf surface while minimizing interference with primary metabolic processes in mesophyll and bundle sheath cells. In semi-arid environments, where plants experience fluctuating herbivory pressure alongside episodic drought, this organization may further enable trichomes to function as sites for both defense and temporary storage of nitrogen and carbon. This hypothesis will require further future inquiry.

### Bulliform cells across development: anatomical persistence and challenges in molecular resolution

Bulliform cells are classically associated with the adult phase and have long been linked to leaf rolling and water status regulation in grasses. In maize, these cells are absent in juvenile leaves and are considered a hallmark of the juvenile-to-adult phase transition (Matschi et al., 2020; Poethig, 1990). In contrast, our SEM-based observations indicate that bulliform-like cells are present in both juvenile and adult sorghum leaves. We find this observation to be true for both domesticated and wild sorghum ecotypes **(Figure 4)**. These findings suggest that, in sorghum, these cell types may be established earlier in development than previously appreciated. In fact, the presence of bulliform cells in juvenile leaves raises the possibility that their functional roles may not be restricted to later developmental stages only. Given their well-established association with leaf rolling and tissue dehydration responses (Matschi et al., 2020), their early presence may contribute to the regulation of leaf water status throughout the sorghum plant life cycle, rather than being confined to adult-phase adaptation. Future studies will be needed to determine the extent to which bulliform cells at juvenile stages are functionally active in leaf hydraulics and water regulation.

Despite their clear anatomical definition, resolving bulliform cells at the molecular level remains challenging. In our snRNA-seq analysis, we were unable to confidently assign a distinct bulliform cell cluster, likely reflecting the current lack of well-validated, cell-type-specific marker genes for this cell type. Although several genes, including *OsZHD1*, *Hal1*, *RL14*, and *ROC5*, have been implicated in bulliform cell development and morphology (Xu et al., 2018), their expression is not restricted to bulliform cells, limiting their utility as definitive markers. Similarly, *ACL1* has been proposed as a candidate marker (Li et al., 2010; Marand et al., 2021), but its expression is also not exclusive, further complicating cell-type resolution using these transcripts as marker genes. This limitation highlights a broader gap in our understanding of bulliform cell biology. The development of robust and specific molecular markers will be essential for resolving these cells in single cell and single nucleus datasets and for dissecting their roles in leaf physiology. In this context, emerging approaches such as spatial transcriptomics offer promising avenues for linking anatomical features with gene expression. Although such methods have been successfully applied in plant tissues such as in wheat spikes (Long et al., 2026; X. Zhang et al., 2026), wheat seeds (Millsteed et al., 2025), and barley grains (Peirats-Llobet et al., 2023), achieving high-resolution spatial transcriptomic profiling in matured C_4_ grass leaves would be particularly valuable for resolving cell types like bulliform cells, which are defined as much by their position and morphology as by their transcriptional signatures. Such advances could provide a foundation for more targeted manipulation of bulliform cell traits beyond leaf rolling, with potential implications for improving leaf water regulation and crop resilience.

### Development overrides cell identity to shape the sorghum leaf transcriptome

Our data indicate that developmental stage, rather than cell identity, is a primary axis of transcriptional organization in sorghum leaves. Specifically, we found that cell types clustered more strongly by juvenile versus adult state than by their canonical cell identities **(Figure 5)**, suggesting that developmental progression imposes a coordinated transcriptional program across diverse cell populations. A similar collapse of cell type-specific expression has been reported under drought, where stress-driven transcriptional responses override intrinsic cell identity (Stata et al., 2025). Consistent with this, recent work in *Arabidopsis* has shown that drought stress can advance transcriptional programs associated with leaf aging, shifting gene expression patterns toward more mature states in a dose-dependent manner, with particularly strong transcriptional shifts observed in mesophyll cells where these changes correlate with reduced leaf growth (Swift et al., 2026). Interestingly, we observe that stage-associated transcriptional signals in our system are also most prominent in mesophyll cells, suggesting that this cell type may play a central role in large-scale transcriptional reprogramming across both stress and developmental contexts in plant leaves. This observation will need to be extended to additional plant species in the future to truly prove this hypothesis. However, our results and other currently available cell level expression results provide initial support for this intriguing hypothesis.

#### Trichomes as multifunctional interfaces for defense and water regulation in sorghum leaves

Trichomes are classically described as epidermal structures primarily associated with biotic defense, including roles in herbivore deterrence and pathogen protection (X. Wang et al., 2021; War et al., 2012). Consistent with this, our analysis revealed that trichome cells in sorghum are enriched for genes associated with secondary metabolism, including the dhurrin biosynthetic pathway **(Figure 3)**. This observation aligns with the known defensive role of dhurrin as a cyanogenic glucoside (Cowan et al., 2022; Darbani et al., 2016) and is further supported by the predominance of glandular trichomes observed in our anatomical analyses **(Figures 2D-E)**, which are widely recognized as sites of specialized metabolite production (Glas et al., 2012; Kennedy, 2003; Shepherd et al., 2005).

In addition to this established defensive role, our data revealed enrichment of multiple aquaporin genes within trichome cells, particularly members of the PIP family that are typically associated with plasma membrane water transport. The co-occurrence of aquaporin expression **(Figure 6C)** with secondary metabolic pathways (**Figure 3A-C)** suggests that trichomes may not function solely as defensive structures but may also participate in local water regulation. While our results do not directly demonstrate functional coupling between these processes, it is possible that aquaporins support the metabolic demands of these cells by facilitating regulated water movement required for metabolite synthesis, storage, or cellular homeostasis.

In other systems, aquaporins have been shown to influence cellular metabolism indirectly, including links to lipid-associated processes and broader metabolic regulation, highlighting that their roles can extend beyond simple water transport (Bi et al., 2025; M. Wang et al., 2016; Wang et al., 2020). Together, these observations suggest that aquaporins in trichome cells may contribute to maintaining the physiological environment necessary for specialized metabolic (Dhurrin) activity. This interpretation is supported by the broader observation that aquaporin expression is developmentally regulated and cell-type specific **(Figure 6)**. In fact, several aquaporin genes examined in this study showed higher expression in juvenile leaves relative to adult leaves, and their expression patterns varied across cell types. Notably, aquaporin enrichment was not uniform within the family, as NIP 2-1 was preferentially enriched in vascular-associated cells rather than trichomes **(Figure 6B).** This heterogeneity suggests that different aquaporin subfamilies may contribute to distinct aspects of water transport across tissues, with trichomes representing one component of a broader, distributed system.

Although trichome studies have traditionally focused on single functional roles, typically examining either water absorption or defense-related processes such as metal detoxification (Li et al., 2023), increasing evidence suggests that these functions are not mutually exclusive. For example, in *Avicennia marina*, trichomes have been shown to perform these dual roles (H. T. Nguyen, Meir, Sack, et al., 2017; H. T. Nguyen, Meir, Wolfe, et al., 2017; Schaepdryver et al., 2022). In our dataset, the co-enrichment of gene ontology terms associated with both defense and water movement, together with the presence of both glandular and non-glandular trichomes **(Figures 2 and 6)**, is consistent with the possibility that sorghum trichomes may also exhibit multifunctionality, although further validation is required. Additionally, support for this hypothesis from our analyses was the non-quantitative observations from our SEM images that glandular trichomes were consistently predominant across the regions analyzed, with only occasional non-glandular trichomes observed.

Additional support for a potential role in water regulation comes from studies of foliar water absorption (FWA). While root-mediated uptake is the dominant pathway for plant hydration, leaves can absorb water directly from the atmosphere, particularly in arid environments (Berry et al., 2019). Although FWA has been documented across diverse plant groups, the mechanisms underlying this process in grasses remain poorly characterized. Proposed pathways include cuticular and stomatal uptake (Xu et al., 2018), but the contribution of trichomes in grasses has not been well explored. Insights from other systems provide a possible mechanistic framework. In the epiphytic species *Tillandsia ionantha*, foliar trichomes mediate water uptake through a biphasic process involving initial surface capture followed by aquaporin-dependent intracellular transport (Ohrui et al., 2007). In this plant, the PIP2 aquaporin subfamily facilitates water movement into underlying tissues after trichome-mediated capture. While functional evidence for such a mechanism in sorghum is lacking, the enrichment of PIP2 aquaporins in trichome cells observed in this study **(Figure 6)** is consistent with a similar potential capacity for localized water movement in sorghum leaves. Anatomical observations further support this possibility. Specifically, in our study, trichomes were consistently found near bulliform cells and, in some cases, positioned directly above them **(Figure 4C)**. This spatial relationship has also been observed in previous anatomical images from maize (Matschi et al., 2020) and sorghum (Hashimoto et al., 2019), suggesting a conserved anatomical arrangement in grasses. Bulliform cells are known to play roles in leaf water status and are characterized by extensive plasmodesmata connections, which could facilitate water redistribution within the leaf (Alvarez et al., 2008). Furthermore, we found that the aquaporin transcript *TIP1-1* was specifically localized to trichome and BC cell regions in our RNAScope validation experiments **(Supplemental Figure 4)**. Together, these findings raise the possibility that trichomes and bulliform cells may form a spatially coordinated system involved in water movement from the leaf surface to internal tissues. This hypothesis should be tested with future research studies.

Previous studies have also proposed that water may move from the trichome base into underlying tissues (Aytasheva et al., 2006), and more recent interpretations have suggested that trichomes may contribute to water delivery to the leaf interior via bulliform cells (Ali et al., 2022). However, these models have largely relied on anatomical observations like ours and lack molecular or functional validation. Additionally, environmental studies have suggested a potential role for trichomes in water acquisition and conservation. For example, trichome density has been shown to increase under dry conditions and decrease with irrigation in *Wigandia urens* (Perez-Estrada et al., 2000), suggesting that trichomes may contribute to adaptation under water-limited conditions. Our findings provide complementary molecular evidence in the form of aquaporin enrichment in trichome cells **(Figure 6 and Supplemental Figure 4)**, although direct functional testing will be necessary to establish whether these proteins actively mediate water transport in this context.

Taken together, our results suggest that sorghum trichomes may function as multifunctional interfaces integrating roles in defense and water regulation. While their enrichment for dhurrin biosynthetic genes supports a well-established role in chemical defense **(Figure 3)**, the concurrent enrichment of aquaporins and their spatial association with bulliform cells raises the possibility that they also contribute to water movement at the leaf surface **(Figure 6 and Supplemental Figure 4)**. Based on these observations, we propose a working model **(Figure 7)** in which trichomes act as initial interfaces for water interaction at the leaf surface, where water from sources such as dew or atmospheric moisture may be captured and subsequently transported across the plasma membrane via aquaporins. This water could then be transferred to bulliform cells (acting as the capacitance (Luo et al., 2021)), which are well positioned to redistribute water within the leaf due to their connectivity and known roles in leaf hydration dynamics. In parallel, aquaporin-mediated regulation of cellular water status within trichomes may support the metabolic processes associated with specialized compound synthesis, including dhurrin production. In this framework, trichomes would not function solely as defensive structures but as integrated sites coordinating surface water interaction, intracellular water regulation, and metabolic activity **(Figure 7)**. While this model remains hypothetical, it provides a conceptual basis for future studies aimed at testing the role of trichomes in foliar water uptake and internal water redistribution in sorghum.

**Figure 7:**
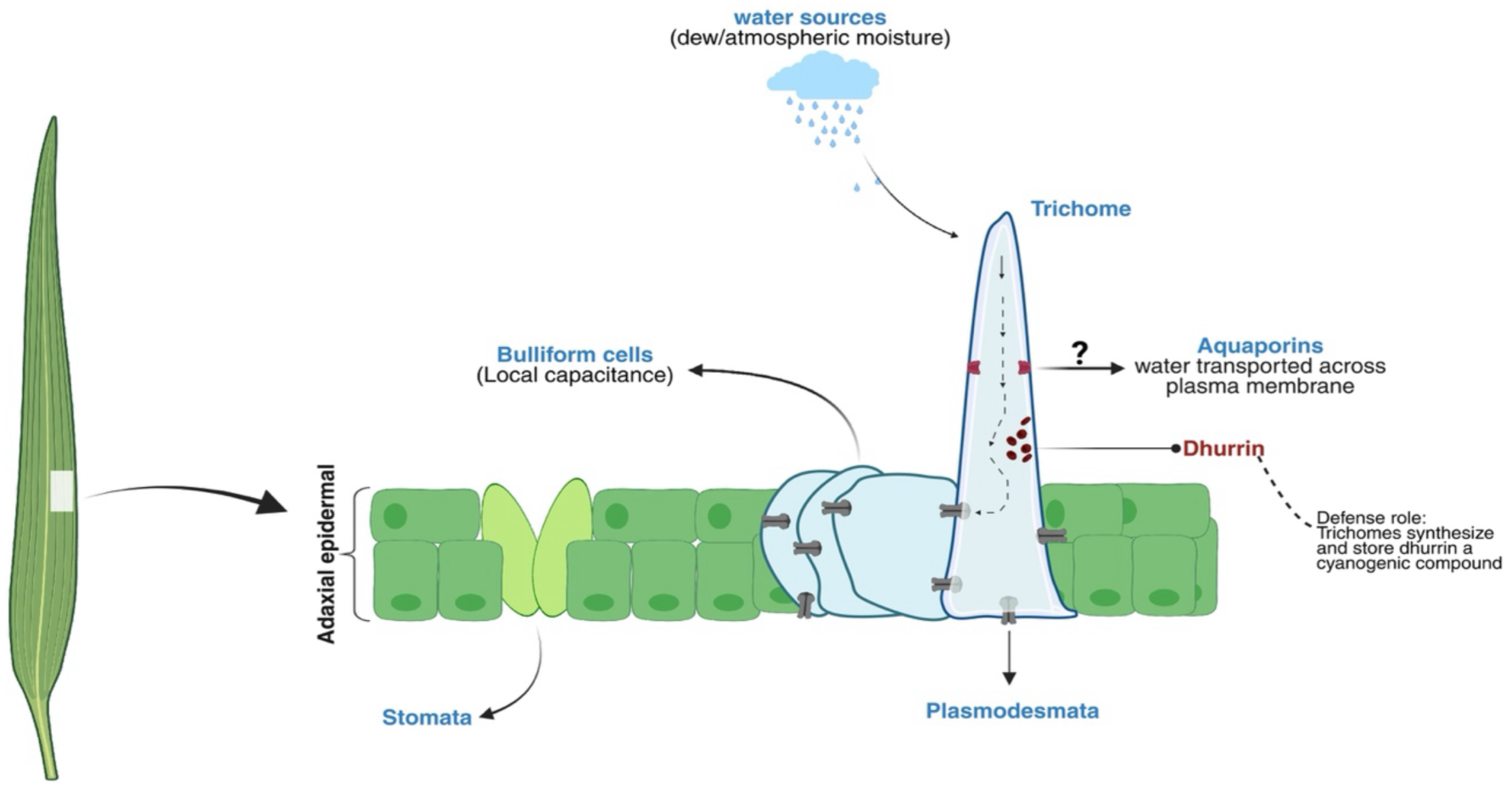
Proposed model for trichome function in water movement and dhurrin biosynthesis. In this model, sorghum trichomes act as epidermal interfaces for surface water uptake via aquaporins and transfer water to adjacent bulliform cells through plasmodesmata, while also contributing to chemical defense via dhurrin biosynthesis.

### Conclusion and implications for sorghum improvement

The results presented in this study demonstrate that trichomes and bulliform cells are present in both juvenile and adult sorghum leaves, suggesting that these epidermal cell types may be established earlier than previously thought. We also observed enrichment of dhurrin biosynthetic genes in trichomes pointing to a possible spatial organization of defense metabolism. While this hypothesis needs further validation, it suggests that targeting specific cell types could be a way to better control dhurrin levels, potentially balancing plant defense with forage safety.

## ACKNOWLEDGMENTS

We thank members of the JK, BH, and BDG labs, both past and present, for helpful discussions. We thank Diep Ganguly for critical feedback and suggestions on the manuscript. This work was funded by NSF grant IOS-1849708 and a grant from University Research Foundation at the University of Pennsylvania to BDG.

## AUTHORS CONTRIBUTIONS

OMA, JK, BH, and BDG conceived and designed the study. OMA performed the snRNA-seq, anatomical, and physiological experiments. OMA, JR, SSB, MP prepared and sequenced the snRNA-seq libraries. OMA, JR, HN, and BDG analyzed the datasets. OMA, JR, HN, AMA, JK, and BDG interpreted the data. OMA and BDG wrote the manuscript. All authors reviewed and edited the manuscript.

## DATA AVAILABILITY

The snRNA-seq data is available through GEO Omnibus under ID GSE311070.

---

**Supplemental Figure 1:**
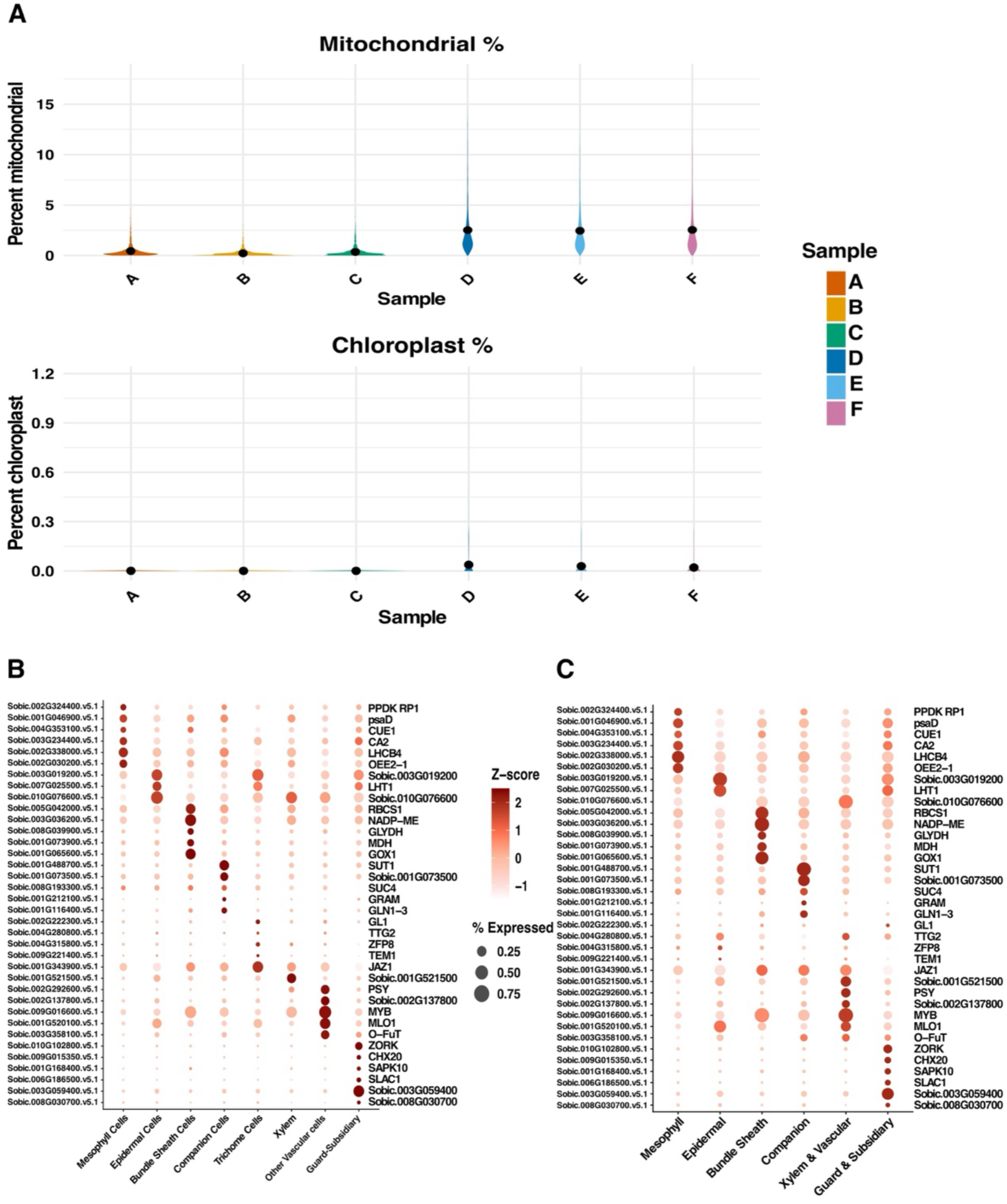
Effect of reference genome choice on marker gene visualization. **(A)** Violin plots of mitochondrial (top) and chloroplast (bottom) transcript percentages across samples A-C uuvenile) and D-F (adult). Overall organellar transcript abundance was low (mitochondrial <∼3%, chloroplast ---0%). (B) Dot plot showing expression of canonical marker genes across cell types using a nuclear-only reference genome. (C) Dot plot showing expression of the same marker genes following alignmenl to a combined reference including mitochondrial (MT) and chloroplast (GP) genomes.

**Supplemental Figure 2:**
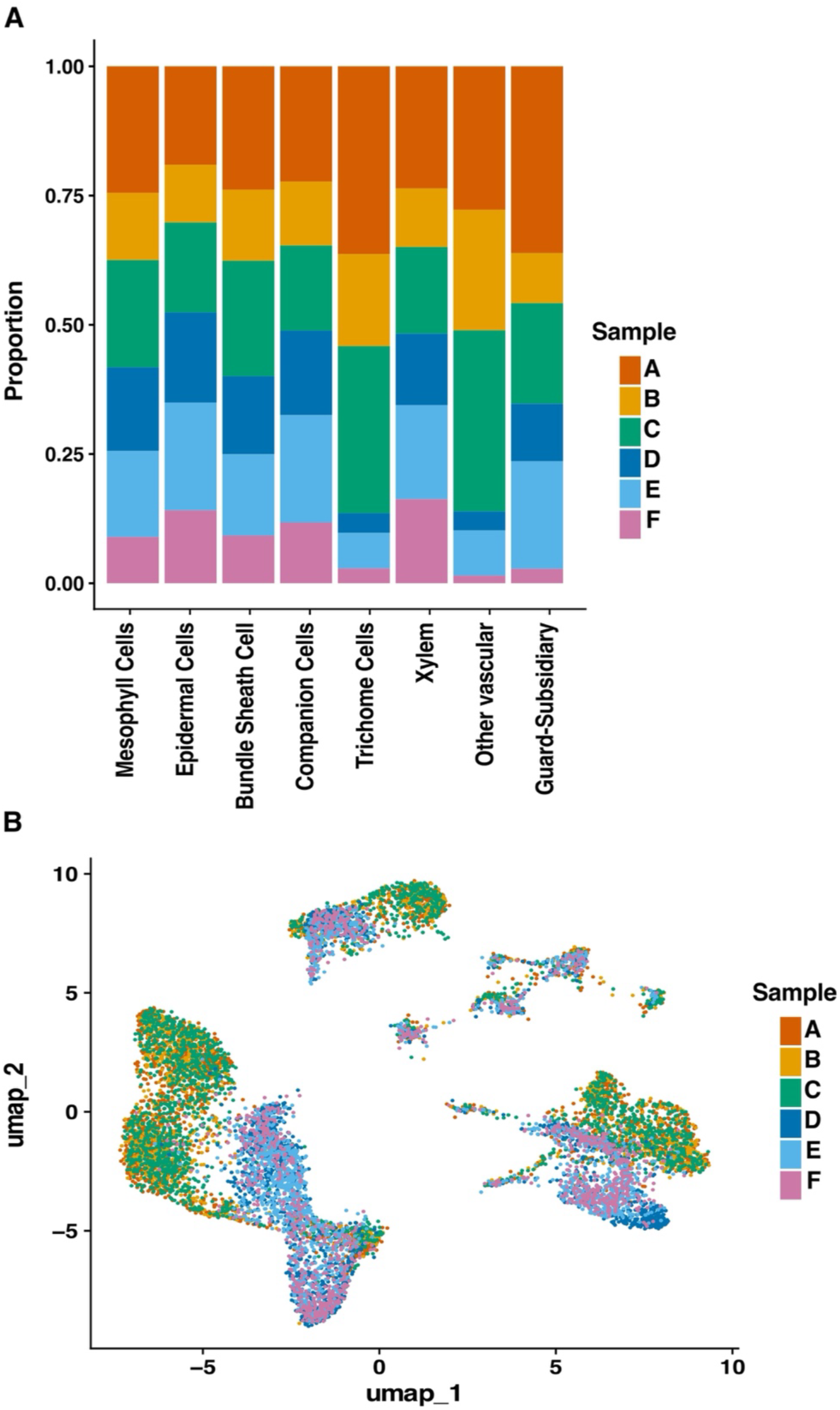
Assessment of technical consistency across pooled snRNA-seq libraries. (A) Proportion of nuclei from each technical channel (A-F) across annotated cell types. Nuclei were isolated from four individual plants per developmental stage, pooled prior to library preparation, and subsequently partitioned into three technical channels (A-C for juvenile and D-F for adult). Each bar represents a cell type, and colors indicate the relative contribution of each channel. Nuclei from multiple channels are represented across cell types, suggesting that clustering is not primarily driven by technical partitioning. (B) UMAP visualization of nuclei colored by technical channel. Nuclei from different channels are broadly distributed across clusters, with no clear channel-specific segregation, supporting overall consistency across libraries

**Supplemental Figure 3:**
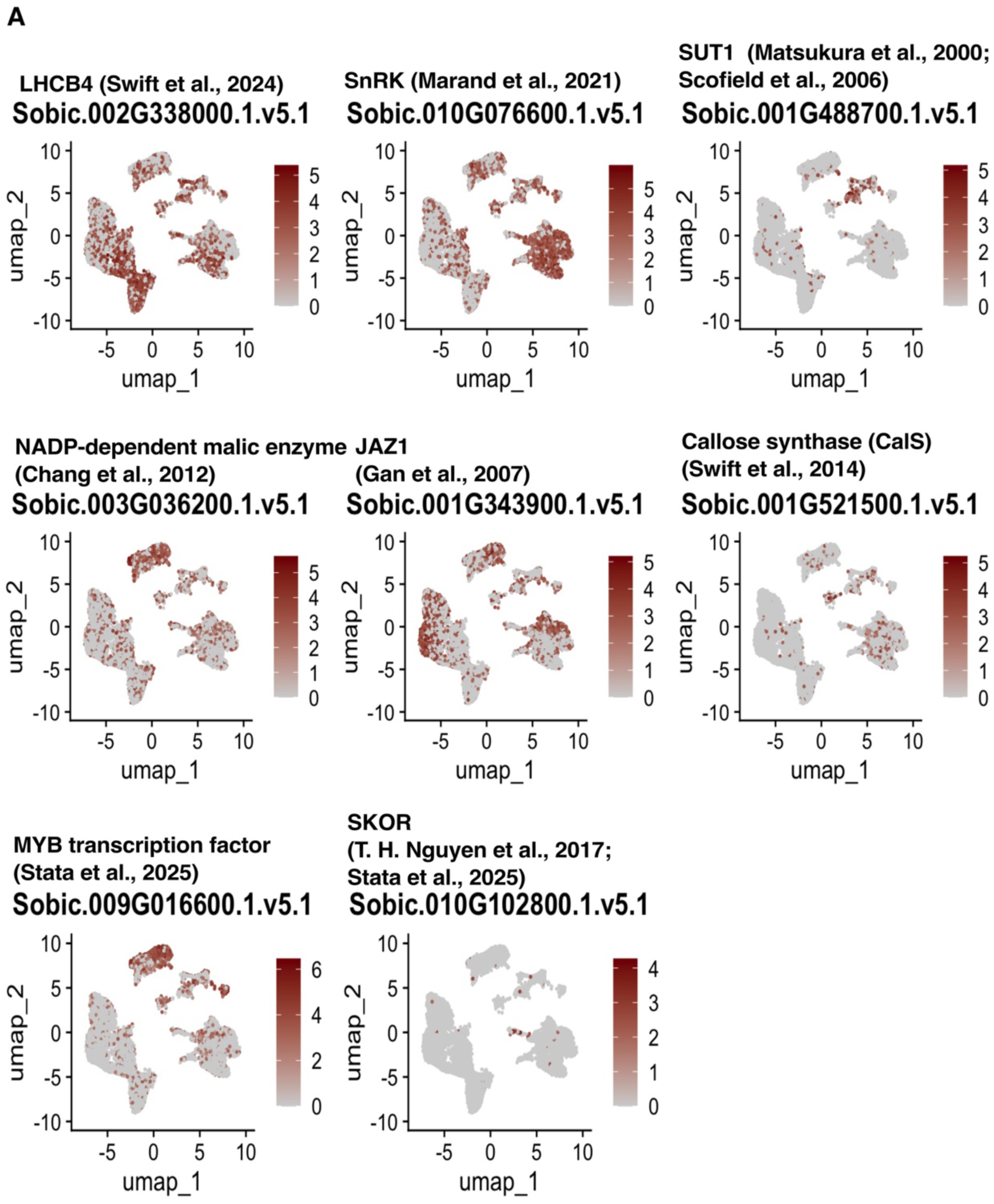
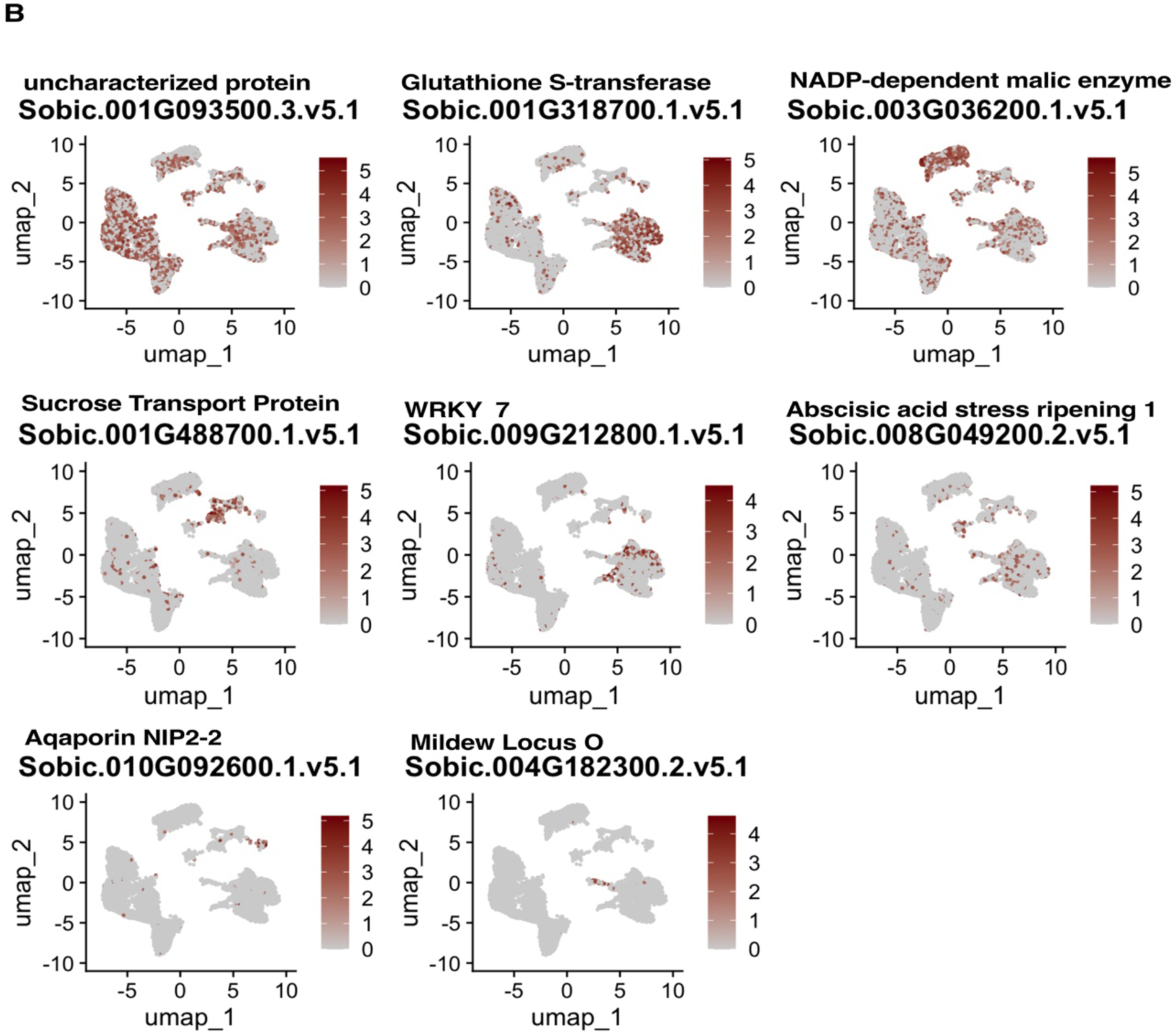
Marker gene expression across sorghum leaf cell types. **(A)** UMAP feature plots showing expression patterns of previously reported marker genes used for cell-type annotation. Color intensity represents normalized expression levels across nuclei. **(B)** UMAP feature plots showing expression patterns of de novo marker genes identified in this study. Color intensity represents normalized expression levels across nuclei.

**Supplemental Figure 4:**
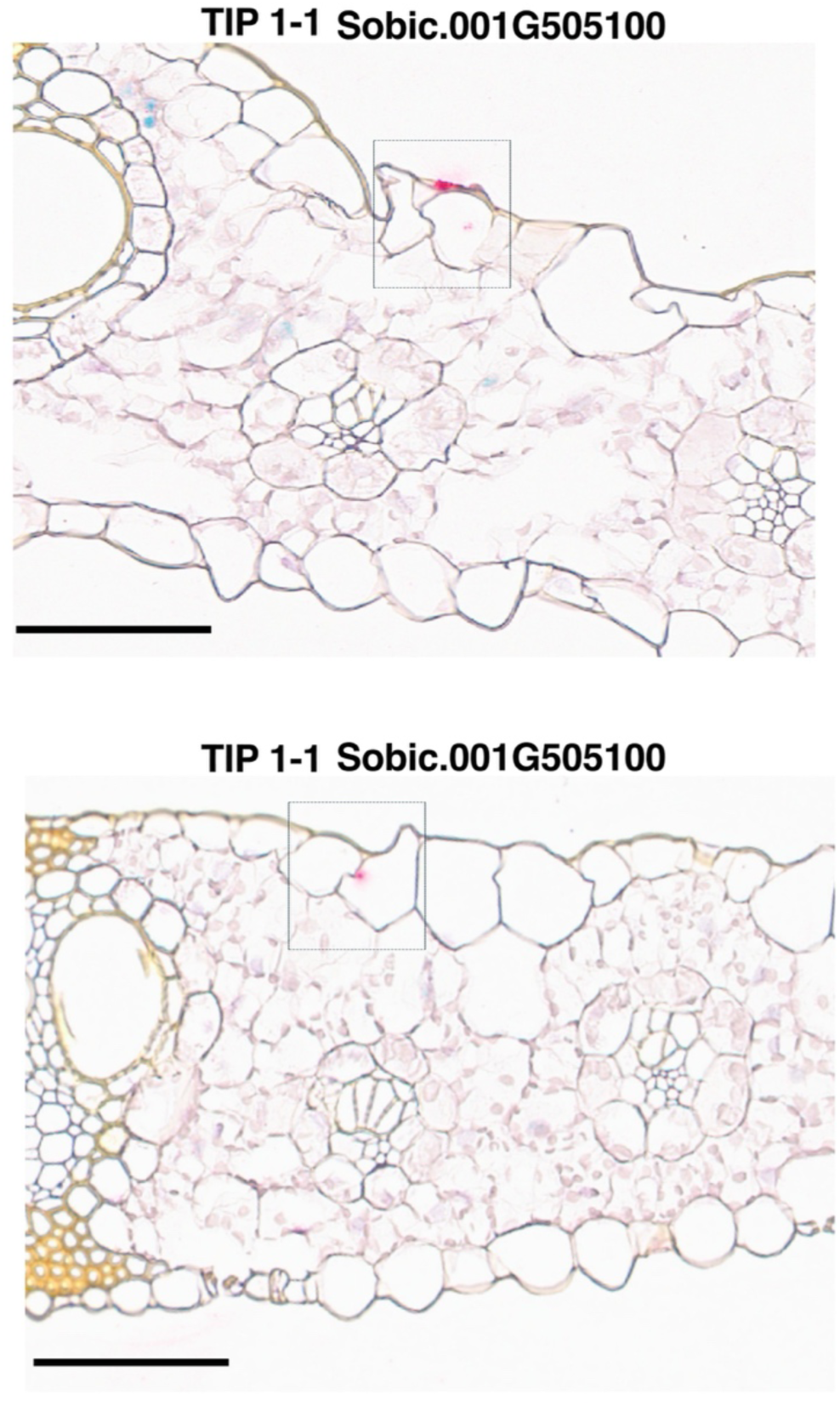
RNAscope localization of TIP1•1 *(Sobic.001G505100).* RNAscope *_in situ_* hybridization of *_T/Pt-1 in_* sorghum leaf cross-sections. Red puncta indicate signal detected primarily in epidermal and sub-epidermal regions, with signal observed in trichome cells (top) and bulliform cells (bottom panel). Scale bars, 20 *µm*.

## SUPPLEMENTAL TABLES

**Supplemental Table 1. Primer sequences and RNAscope probes used in this study.**

**Supplemental Table 2. Gene ontology (GO) biological process enrichment for each cell-type cluster.**

**Supplemental Table 3. Marker genes from published single-cell studies used for cell-type annotation. Curated reference markers compiled from prior plant single-cell/nucleus papers and used to guide initial cluster labels in this study.**

**Supplemental Table 4. All markers identified in this study by cell types.**

**Supplemental Table 5. All markers identified in this study by cell types and development stage.**

**Supplemental Table 6. Differentially expressed genes (DEGs) in trichome cells between juvenile and adult stages.**

## REFERENCES

Afzal, Z., Howton, T., Sun, Y., & Mukhtar, M. (2016). The Roles of Aquaporins in Plant Stress Responses. Journal of Developmental Biology, 4(1), 9. 10.3390/jdb4010009

Ali, Z., Merrium, S., Habib-ur-Rahman, M., Hakeem, S., Saddique, M. A. B., & Sher, M. A. (2022). Wetting mechanism and morphological adaptation; leaf rolling enhancing atmospheric water acquisition in wheat crop—A review. Environmental Science and Pollution Research, 29(21), 30967–30985. 10.1007/s11356-022-18846-3

Al-Salman, Y., Cano, F. J., Pan, L., Koller, F., Piñeiro, J., Jordan, D., & Ghannoum, O. (2023). Anatomical drivers of stomatal conductance in sorghum lines with different leaf widths grown under different temperatures. Plant, Cell & Environment, 46(7), 2142–2158. 10.1111/pce.14592

Alvarez, J. M., Rocha, J. F., & Machado, S. R. (2008). Bulliform cells in Loudetiopsis chrysothrix (Nees) Conert and Tristachya leiostachya Nees (Poaceae): Structure in relation to function. Brazilian Archives of Biology and Technology, 51(1), 113–119. 10.1590/S1516-89132008000100014

Asai, K., Satoh, N., Sasaki, H., Satoh, H., & Nagato, Y. (2002). A rice heterochronic mutant, *mori1*, is defective in the juvenile-adult phase change. Development, 129(1), 265–273. 10.1242/dev.129.1.265

Becraft, P. W., Li, K., Dey, N., & Asuncion-Crabb, Y. (2002). The maize *dek1* gene functions in embryonic pattern formation and cell fate specification. Development, 129(22), 5217–5225. 10.1242/dev.129.22.5217

Berry, Z. C., Emery, N. C., Gotsch, S. G., & Goldsmith, G. R. (2019). Foliar water uptake: Processes, pathways, and integration into plant water budgets. *Plant*, Cell & Environment, 42(2), 410–423. 10.1111/pce.13439

Bi, Y., Pang, S., Liu, Y., Cheng, J., Ma, Q., Song, A., Yin, X., & Wang, J. (2025). Aquaporins in lipid metabolism: Functions and regulation in health and disease. Lipids in Health and Disease, 24(1), 336. 10.1186/s12944-025-02727-y

Bjarnholt, N., Neilson, E. H. J., Crocoll, C., Jørgensen, K., Motawia, M. S., Olsen, C. E., Dixon, D. P., Edwards, R., & Møller, B. L. (2018). Glutathione transferases catalyze recycling of auto-toxic cyanogenic glucosides in sorghum. The Plant Journal, 94(6), 1109–1125. 10.1111/tpj.13923

Bongard-Pierce, D. K., Evans, M. M. S., & Poethig, R. S. (1996). Heteroblastic Features of Leaf Anatomy in Maize and Their Genetic Regulation. International Journal of Plant Sciences, 157(4), 331–340. 10.1086/297353

Byrt, C. S., Zhang, R. Y., Magrath, I., Chan, K. X., De Rosa, A., & McGaughey, S. (2023). Exploring aquaporin functions during changes in leaf water potential. Frontiers in Plant Science, 14, 1213454. 10.3389/fpls.2023.1213454

Cal, A. J., Sanciangco, M., Rebolledo, M. C., Luquet, D., Torres, R. O., McNally, K. L., & Henry, A. (2019). Leaf morphology, rather than plant water status, underlies genetic variation of rice leaf rolling under drought. Plant, Cell & Environment, 42(5), 1532–1544. 10.1111/pce.13514

Chandna, R., Augustine, R., & Bisht, N. C. (2012). Evaluation of Candidate Reference Genes for Gene Expression Normalization in Brassica juncea Using Real Time Quantitative RT-PCR. PLoS ONE, 7(5), e36918. 10.1371/journal.pone.0036918

Chang, Y.-M., Liu, W.-Y., Shih, A. C.-C., Shen, M.-N., Lu, C.-H., Lu, M.-Y. J., Yang, H.-W., Wang, T.-Y., Chen, S. C.-C., Chen, S. M., Li, W.-H., & Ku, M. S. B. (2012). Characterizing Regulatory and Functional Differentiation between Maize Mesophyll and Bundle Sheath Cells by Transcriptomic Analysis. Plant Physiology, 160(1), 165–177. 10.1104/pp.112.203810

Cowan, M., Møller, B. L., Norton, S., Knudsen, C., Crocoll, C., Furtado, A., Henry, R., Blomstedt, C., & Gleadow, R. M. (2022). Cyanogenesis in the Sorghum Genus: From Genotype to Phenotype. Genes, 13(1), 140. 10.3390/genes13010140

Darbani, B., Motawia, M. S., Olsen, C. E., Nour-Eldin, H. H., Møller, B. L., & Rook, F. (2016). The biosynthetic gene cluster for the cyanogenic glucoside dhurrin in Sorghum bicolor contains its co-expressed vacuolar MATE transporter. Scientific Reports, 6(1), 37079. 10.1038/srep37079

Denyer, T., Ma, X., Klesen, S., Scacchi, E., Nieselt, K., & Timmermans, M. C. P. (2019). Spatiotemporal Developmental Trajectories in the Arabidopsis Root Revealed Using High-Throughput Single-Cell RNA Sequencing. Developmental Cell, 48(6), 840–852.e5. 10.1016/j.devcel.2019.02.022

Develey-Rivière, M., & Galiana, E. (2007). Resistance to pathogens and host developmental stage: A multifaceted relationship within the plant kingdom. New Phytologist, 175(3), 405–416. 10.1111/j.1469-8137.2007.02130.x

Doody, E., Zha, Y., He, J., & Poethig, R. S. (2022). The genetic basis of natural variation in the timing of vegetative phase change in *Arabidopsis thaliana*. Development, 149(10), dev200321. 10.1242/dev.200321

Döring, F., Streubel, M., Bräutigam, A., & Gowik, U. (2016). Most photorespiratory genes are preferentially expressed in the bundle sheath cells of the C_4_ grass *Sorghum bicolor*. Journal of Experimental Botany, 67(10), 3053–3064. 10.1093/jxb/erw041

Gan, Y., Liu, C., Yu, H., & Broun, P. (2007). Integration of cytokinin and gibberellin signalling by *Arabidopsis* transcription factors GIS, ZFP8 and GIS2 in the regulation of epidermal cell fate. Development, 134(11), 2073–2081. 10.1242/dev.005017

Gimeno, J., Eattock, N., Van Deynze, A., & Blumwald, E. (2014). Selection and Validation of Reference Genes for Gene Expression Analysis in Switchgrass (Panicum virgatum) Using Quantitative Real-Time RT-PCR. PLoS ONE, 9(3), e91474. 10.1371/journal.pone.0091474

Glas, J., Schimmel, B., Alba, J., Escobar-Bravo, R., Schuurink, R., & Kant, M. (2012). Plant Glandular Trichomes as Targets for Breeding or Engineering of Resistance to Herbivores. International Journal of Molecular Sciences, 13(12), 17077–17103. 10.3390/ijms131217077

Grones, C., Eekhout, T., Shi, D., Neumann, M., Berg, L. S., Ke, Y., Shahan, R., Cox, K. L., Gomez-Cano, F., Nelissen, H., Lohmann, J. U., Giacomello, S., Martin, O. C., Cole, B., Wang, J.-W., Kaufmann, K., Raissig, M. T., Palfalvi, G., Greb, T., … De Rybel, B. (2024). Best practices for the execution, analysis, and data storage of plant single-cell/nucleus transcriptomics. The Plant Cell, 36(4), 812–828. 10.1093/plcell/koae003

Guillotin, B., & D. Birnbaum, K. (2025). Plant nuclei extraction protocol for single nuclei RNA-seq v1. 10.17504/protocols.io.yxmvmb3d6g3p/v1

Guillotin, B., Rahni, R., Passalacqua, M., Mohammed, M. A., Xu, X., Raju, S. K., Ramírez, C. O., Jackson, D., Groen, S. C., Gillis, J., & Birnbaum, K. D. (2023). A pan-grass transcriptome reveals patterns of cellular divergence in crops. Nature, 617(7962), 785–791. 10.1038/s41586-023-06053-0

Hashimoto, S., Tezuka, T., & Yokoi, S. (2019). Morphological changes during juvenile-to-adult phase transition in sorghum. Planta, 250(5), 1557–1566. 10.1007/s00425-019-03251-x

Hill, R. D., Robertson, S. M., Igamberdiev, A. U., Mira, M. M., Wilkins, O., & Stasolla, C. (2026). Discrete and cell-specific hypoxic responses in Arabidopsis roots resolved by single-nuclei transcriptomics. New Phytologist, 249(6), 2652–2667. 10.1111/nph.70874

Hu, L., & Yang, L. (2019). Time to Fight: Molecular Mechanisms of Age-Related Resistance. Phytopathology®, 109(9), 1500–1508. 10.1094/PHYTO-11-18-0443-RVW

Huijser, P., & Schmid, M. (2011). The control of developmental phase transitions in plants. Development, 138(19), 4117–4129. 10.1242/dev.063511

Illouz-Eliaz, N., Yu, J., Swift, J., Lande, K., Jow, B., Partida-Garcia, L., Tuang, Z. K., Lee, T. A., Yaaran, A., Gomez-Castanon, R., Owens, W., Bowman, C. R., Osgood, E., Nery, J. R., Nobori, T., Zait, Y., Burdman, S., & Ecker, J. R. (2025). Drought recovery in plants triggers a cell-state-specific immune activation. Nature Communications, 16(1), 8095. 10.1038/s41467-025-63467-2

Jean-Baptiste, K., McFaline-Figueroa, J. L., Alexandre, C. M., Dorrity, M. W., Saunders, L., Bubb, K. L., Trapnell, C., Fields, S., Queitsch, C., & Cuperus, J. T. (2019). Dynamics of Gene Expression in Single Root Cells of *Arabidopsis thaliana*. The Plant Cell, 31(5), 993–1011. 10.1105/tpc.18.00785

Kennedy, G. G. (2003). Tomato, Pests, Parasitoids, and Predators: Tritrophic Interactions Involving the Genus *Lycopersicon*. Annual Review of Entomology, 48(1), 51–72. 10.1146/annurev.ento.48.091801.112733

Kim, J.-Y., Symeonidi, E., Pang, T. Y., Denyer, T., Weidauer, D., Bezrutczyk, M., Miras, M., Zöllner, N., Hartwig, T., Wudick, M. M., Lercher, M., Chen, L.-Q., Timmermans, M. C. P., & Frommer, W. B. (2021). Distinct identities of leaf phloem cells revealed by single cell transcriptomics. The Plant Cell, 33(3), 511–530. 10.1093/plcell/koaa060

Koleva, D. T., Liu, M., Dusak, B., Ghosh, S., Krogh, C. T., Hellebek, I. R., Cortsen, M. T., Motawie, M. S., Jørgensen, F. S., McKinley, B. A., Mullet, J. E., Sørensen, M., & Møller, B. L. (2025). Amino acid substrate specificities and tissue expression profiles of the nine CYP79A encoding genes in *Sorghum bicolor*. Physiologia Plantarum, 177(1), e70029. 10.1111/ppl.70029

Larkin, J. C., Oppenheimer, D. G., Lloyd, A. M., Paparozzi, E. T., & Marks, M. D. (1994). Roles of the GLABROUS1 and TRANSPARENT TESTA GLABRA Genes in Arabidopsis Trichome Development. The Plant Cell, 1065–1076. 10.1105/tpc.6.8.1065

Lawson, E. J. R., & Poethig, R. S. (1995). Shoot development in plants: Time for a change. Trends in Genetics, 11(7), 263–268. 10.1016/S0168-9525(00)89072-1

Lee, T. A., Nobori, T., Illouz-Eliaz, N., Xu, J., Jow, B., Nery, J. R., & Ecker, J. R. (2023). A Single-Nucleus Atlas of Seed-to-Seed Development in Arabidopsis. Plant Biology. 10.1101/2023.03.23.533992

Li, C., Mo, Y., Wang, N., Xing, L., Qu, Y., Chen, Y., Yuan, Z., Ali, A., Qi, J., Fernández, V., Wang, Y., & Kopittke, P. M. (2023). The overlooked functions of trichomes: Water absorption and metal detoxication. Plant, Cell & Environment, 46(3), 669–687. 10.1111/pce.14530

Li, L., Shi, Z.-Y., Li, L., Shen, G.-Z., Wang, X.-Q., An, L.-S., & Zhang, J.-L. (2010). Overexpression of ACL1 (abaxially curled leaf 1) Increased Bulliform Cells and Induced Abaxial Curling of Leaf Blades in Rice. Molecular Plant, 3(5), 807–817. 10.1093/mp/ssq022

Li, Y., Wen, S., Li, Z., Liu, R., Zhang, Z., Li, Y., Lyu, D., & Jian, H. (2025). The evolution of the aquaporin gene family and drought tolerance mechanisms in green plants. Horticulture Research, 12(11), uhaf209. 10.1093/hr/uhaf209

Liu, Y., Yang, M., Deng, Y., Su, G., Enninful, A., Guo, C. C., Tebaldi, T., Zhang, D., Kim, D., Bai, Z., Norris, E., Pan, A., Li, J., Xiao, Y., Halene, S., & Fan, R. (2020). High-Spatial-Resolution Multi-Omics Sequencing via Deterministic Barcoding in Tissue. Cell, 183(6), 1665–1681.e18. 10.1016/j.cell.2020.10.026

Long, K. A., Lister, A., Jones, M. R. W., Adamski, N. M., Ellis, R. E., Chedid, C., Carpenter, S. J., Liu, X., Backhaus, A. E., Goldson, A., Knitlhoffer, V., Pei, Y., Vickers, M., Steuernagel, B., Kaithakottil, G. G., Xiao, J., Haerty, W., Macaulay, I. C., & Uauy, C. (2026). Spatial transcriptomics reveals expression gradients in developing wheat inflorescences at cellular resolution. The Plant Cell, 38(1), koaf282. 10.1093/plcell/koaf282

Luo, Y., Ho, C.-L., Helliker, B. R., & Katifori, E. (2021). Leaf Water Storage and Robustness to Intermittent Drought: A Spatially Explicit Capacitive Model for Leaf Hydraulics. Frontiers in Plant Science, 12, 725995. 10.3389/fpls.2021.725995

Marand, A. P., Chen, Z., Gallavotti, A., & Schmitz, R. J. (2021). A cis-regulatory atlas in maize at single-cell resolution. Cell, 184(11), 3041–3055.e21. 10.1016/j.cell.2021.04.014

Matías-Hernández, L., Aguilar-Jaramillo, A. E., Osnato, M., Weinstain, R., Shani, E., Suárez-López, P., & Pelaz, S. (2016). TEMPRANILLO Reveals the Mesophyll as Crucial for Epidermal Trichome Formation. Plant Physiology, 170(3), 1624–1639. 10.1104/pp.15.01309

Matschi, S., Vasquez, M. F., Bourgault, R., Steinbach, P., Chamness, J., Kaczmar, N., Gore, M. A., Molina, I., & Smith, L. G. (2020). Structure-function analysis of the maize bulliform cell cuticle and its potential role in dehydration and leaf rolling. Plant Direct, 4(10), e00282. 10.1002/pld3.282

Matsukura, C., Saitoh, T., Hirose, T., Ohsugi, R., Perata, P., & Yamaguchi, J. (2000). Sugar Uptake and Transport in Rice Embryo. Expression of Companion Cell-Specific Sucrose Transporter (*OsSUT1*) Induced by Sugar and Light. Plant Physiology, 124(1), 85–94. 10.1104/pp.124.1.85

Mendieta, J. P., Tu, X., Jiang, D., Yan, H., Zhang, X., Marand, A. P., Zhong, S., & Schmitz, R. J. (2024). Investigating the *cis-* regulatory basis of C_3_ and C_4_ photosynthesis in grasses at single-cell resolution. Proceedings of the National Academy of Sciences, 121(40), e2402781121. 10.1073/pnas.2402781121

Millsteed, T., Kainer, D., Sullivan, R., Sun, X., Li, K. L., Mao, L., Macdonald, A., & Henry, R. J. (2025). Spatial Transcriptomics of Developing Wheat Seed Reveals Concentric Gene Expression Zones and Subgenome Biased Expression of Key Genes. Plant Biotechnology Journal, 23(12), 5934–5949. 10.1111/pbi.70351

Min, M. K., Choi, E.-H., Kim, J.-A., Yoon, I. S., Han, S., Lee, Y., Lee, S., & Kim, B.-G. (2019). Two Clade A Phosphatase 2Cs Expressed in Guard Cells Physically Interact With Abscisic Acid Signaling Components to Induce Stomatal Closure in Rice. Rice, 12(1), 37. 10.1186/s12284-019-0297-7

Morris, G. P., Harder, A. M., Healey, A. L., McLaughlin, C. M., Rifkin, J. L., Cruet-Burgos, C., Jenkins, J. W., Shu, S., Spiekerman, J. J., VanGessel, C. J., Agnew, E., Audebert, A., Barry, K., Baxter, I., Beurier, G., Boston, L. B., Boyles, R. E., Brady, S. M., Bunting, V., … Lovell, J. T. (2026). A sorghum pangenome reference improves global crop trait discovery. Nature. 10.1038/s41586-026-10229-9

Moulia, B. (2000). Leaves as Shell Structures: Double Curvature, Auto-Stresses, and Minimal Mechanical Energy Constraints on Leaf Rolling in Grasses. Journal of Plant Growth Regulation, 19(1), 19–30. 10.1007/s003440000004

Mundia, C. W., Secchi, S., Akamani, K., & Wang, G. (2019). A Regional Comparison of Factors Affecting Global Sorghum Production: The Case of North America, Asia and Africa’s Sahel. Sustainability, 11(7), 2135. 10.3390/su11072135

Nguyen, H. T., Meir, P., Sack, L., Evans, J. R., Oliveira, R. S., & Ball, M. C. (2017). Leaf water storage increases with salinity and aridity in the mangrove *Avicennia marina*: Integration of leaf structure, osmotic adjustment and access to multiple water sources. *Plant*, Cell & Environment, 40(8), 1576–1591. 10.1111/pce.12962

Nguyen, H. T., Meir, P., Wolfe, J., Mencuccini, M., & Ball, M. C. (2017). Plumbing the depths: Extracellular water storage in specialized leaf structures and its functional expression in a three-domain pressure –volume relationship. Plant, Cell & Environment, 40(7), 1021–1038. 10.1111/pce.12788

Nguyen, T. H., Huang, S., Meynard, D., Chaine, C., Michel, R., Roelfsema, M. R. G., Guiderdoni, E., Sentenac, H., & Véry, A.-A. (2017). A Dual Role for the OsK5.2 Ion Channel in Stomatal Movements and K^+^ Loading into Xylem Sap. Plant Physiology, 174(4), 2409–2418. 10.1104/pp.17.00691

Ohrui, T., Nobira, H., Sakata, Y., Taji, T., Yamamoto, C., Nishida, K., Yamakawa, T., Sasuga, Y., Yaguchi, Y., Takenaga, H., & Tanaka, S. (2007). Foliar trichome- and aquaporin-aided water uptake in a drought-resistant epiphyte Tillandsia ionantha Planchon. Planta, 227(1), 47–56. 10.1007/s00425-007-0593-0

Okamoto, S., Negishi, K., Toyama, Y., Ushijima, T., & Morohashi, K. (2020). Leaf Trichome Distribution Pattern in Arabidopsis Reveals Gene Expression Variation Associated with Environmental Adaptation. Plants, 9(7), 909. 10.3390/plants9070909

Orkwiszewski, J. A. J., & Poethig, R. S. (2000). Phase identity of the maize leaf is determined after leaf initiation. Proceedings of the National Academy of Sciences, 97(19), 10631–10636. 10.1073/pnas.180301597

Peirats-Llobet, M., Yi, C., Liew, L. C., Berkowitz, O., Narsai, R., Lewsey, M. G., & Whelan, J. (2023). Spatially resolved transcriptomic analysis of the germinating barley grain. Nucleic Acids Research, 51(15), 7798–7819. 10.1093/nar/gkad521

Perez-Estrada, L. B., Cano-Santana, Z., & Oyama, K. (2000). Variation in leaf trichomes of Wigandia urens: Environmental factors and physiological consequences. Tree Physiology, 20(9), 629–632. 10.1093/treephys/20.9.629

Poethig, R. S. (1990). Phase Change and the Regulation of Shoot Morphogenesis in Plants. Science, 250(4983), 923–930. 10.1126/science.250.4983.923

Poethig, R. S. (2003). Phase Change and the Regulation of Developmental Timing in Plants. Science, 301(5631), 334–336. 10.1126/science.1085328

Poethig, R. S. (2013). Vegetative Phase Change and Shoot Maturation in Plants. In Current Topics in Developmental Biology (Vol. 105, pp. 125–152). Elsevier. 10.1016/B978-0-12-396968-2.00005-1

Poethig, R. S., & Fouracre, J. (2024). Temporal regulation of vegetative phase change in plants. Developmental Cell, 59(1), 4–19. 10.1016/j.devcel.2023.11.010

Qi, T., Song, S., Ren, Q., Wu, D., Huang, H., Chen, Y., Fan, M., Peng, W., Ren, C., & Xie, D. (2011). The Jasmonate-ZIM-Domain Proteins Interact with the WD-Repeat/bHLH/MYB Complexes to Regulate Jasmonate-Mediated Anthocyanin Accumulation and Trichome Initiation in *Arabidopsis thaliana*. The Plant Cell, 23(5), 1795–1814. 10.1105/tpc.111.083261

Rankenberg, T., Geldhof, B., Van Veen, H., Holsteens, K., Van De Poel, B., & Sasidharan, R. (2021). Age-Dependent Abiotic Stress Resilience in Plants. Trends in Plant Science, 26(7), 692–705. 10.1016/j.tplants.2020.12.016

Reddy, P. S., Rao, T. S. R. B., Sharma, K. K., & Vadez, V. (2015). Genome-wide identification and characterization of the aquaporin gene family in Sorghum bicolor (L.). Plant Gene, 1, 18–28. 10.1016/j.plgene.2014.12.002

Ryu, K. H., Huang, L., Kang, H. M., & Schiefelbein, J. (2019). Single-Cell RNA Sequencing Resolves Molecular Relationships Among Individual Plant Cells. Plant Physiology, 179(4), 1444–1456. 10.1104/pp.18.01482

Schaepdryver, K. H. D., Goossens, W., Naseef, A., Kalpuzha Ashtamoorthy, S., & Steppe, K. (2022). Foliar Water Uptake Capacity in Six Mangrove Species. Forests, 13(6), 951. 10.3390/f13060951

Schreel, J. D. M., Leroux, O., Goossens, W., Brodersen, C., Rubinstein, A., & Steppe, K. (2020). Identifying the pathways for foliar water uptake in beech (*Fagus sylvatica* L.): A major role for trichomes. The Plant Journal, 103(2), 769–780. 10.1111/tpj.14770

Scofield, G. N., Aoki, N., Hirose, T., Takano, M., Jenkins, C. L. D., & Furbank, R. T. (2006). The role of the sucrose transporter, OsSUT1, in germination and early seedling growth and development of rice plants. Journal of Experimental Botany, 58(3), 483–495. 10.1093/jxb/erl217

Serrano-Ron, L., Perez-Garcia, P., Sanchez-Corrionero, A., Gude, I., Cabrera, J., Ip, P.-L., Birnbaum, K. D., & Moreno-Risueno, M. A. (2021). Reconstruction of lateral root formation through single-cell RNA sequencing reveals order of tissue initiation. Molecular Plant, 14(8), 1362–1378. 10.1016/j.molp.2021.05.028

Shepherd, R. W., Bass, W. T., Houtz, R. L., & Wagner, G. J. (2005). Phylloplanins of Tobacco Are Defensive Proteins Deployed on Aerial Surfaces by Short Glandular Trichomes. The Plant Cell, 17(6), 1851–1861. 10.1105/tpc.105.031559

Shulse, C. N., Cole, B. J., Ciobanu, D., Lin, J., Yoshinaga, Y., Gouran, M., Turco, G. M., Zhu, Y., O’Malley, R. C., Brady, S. M., & Dickel, D. E. (2019). High-Throughput Single-Cell Transcriptome Profiling of Plant Cell Types. Cell Reports, 27(7), 2241–2247.e4. 10.1016/j.celrep.2019.04.054

Song, Q., Ando, A., Jiang, N., Ikeda, Y., & Chen, Z. J. (2020). Single-cell RNA-seq analysis reveals ploidy-dependent and cell-specific transcriptome changes in Arabidopsis female gametophytes. Genome Biology, 21(1), 178. 10.1186/s13059-020-02094-0

Stata, M., Greenblum, S., Karia, P., Koriabine, M., Yoshinaga, Y., Keymanesh, K., Zhao, C., O’Malley, R. C., & Rhee, S. Y. (2025). *Single-cell-level response to drought in* Sorghum bicolor *reveals novel targets for improving water use efficiency*. Plant Biology. 10.1101/2025.08.28.671794

Strable, J., Borsuk, L., Nettleton, D., Schnable, P. S., & Irish, E. E. (2008). Microarray analysis of vegetative phase change in maize. The Plant Journal, 56(6), 1045–1057. 10.1111/j.1365-313X.2008.03661.x

Sun, G., Xia, M., Li, J., Ma, W., Li, Q., Xie, J., Bai, S., Fang, S., Sun, T., Feng, X., Guo, G., Niu, Y., Hou, J., Ye, W., Ma, J., Guo, S., Wang, H., Long, Y., Zhang, X., … Song, C.-P. (2022). The maize single-nucleus transcriptome comprehensively describes signaling networks governing movement and development of grass stomata. *The Plant Cell*, koac047. 10.1093/plcell/koac047

Swift, J., Luginbuehl, L. H., Hua, L., Schreier, T. B., Donald, R. M., Stanley, S., Wang, N., Lee, T. A., Nery, J. R., Ecker, J. R., & Hibberd, J. M. (2024). Exaptation of ancestral cell-identity networks enables C4 photosynthesis. Nature, 636(8041), 143–150. 10.1038/s41586-024-08204-3

Swift, J., Wu, X., Xu, J., Procko, C., Jain, T., Illouz-Eliaz, N., Nery, J. R., Chory, J., & Ecker, J. R. (2026). Stress drives plasticity in leaf ageing transcriptional dynamics in Arabidopsis thaliana. Nature Plants. 10.1038/s41477-026-02254-3

Sylvester, A. W., & Smith, L. G. (2009). Cell Biology of Maize Leaf Development. In J. L. Bennetzen & S. C. Hake (Eds.), Handbook of Maize: Its Biology (pp. 179–203). Springer New York. 10.1007/978-0-387-79418-1_10

Telfer, A., Bollman, K. M., & Poethig, R. S. (1997). Phase change and the regulation of trichome distribution in *Arabidopsis thaliana*. Development, 124(3), 645–654. 10.1242/dev.124.3.645

Tenorio Berrío, R., Verhelst, E., Eekhout, T., Grones, C., De Veylder, L., De Rybel, B., & Dubois, M. (2025). Dual and spatially resolved drought responses in the Arabidopsis leaf mesophyll revealed by single-cell transcriptomics. New Phytologist, 246(3), 840–858. 10.1111/nph.20446

Thayer, S. S., & Conn, E. E. (1981). Subcellular Localization of Dhurrin β-Glucosidase and Hydroxynitrile Lyase in the Mesophyll Cells of *Sorghum* Leaf Blades. Plant Physiology, 67(4), 617–622. 10.1104/pp.67.4.617

Triplett, E., Hayes, C., Emendack, Y., Longing, S., Monclova, C., Simpson, C., & Laza, H. E. (2023). Leaf structural traits mediating pre-existing physical innate resistance to sorghum aphid in sorghum under uninfested conditions. Planta, 258(2), 46. 10.1007/s00425-023-04194-0

Truernit, E., & Sauer, N. (1995). The promoter of the Arabidopsis thaliana SUC2 sucrose-H+ symporter gene directs expression of ?-glucuronidase to the phloem: Evidence for phloem loading and unloading by SUC2. Planta, 196(3). 10.1007/BF00203657

Turco, G. M., Rodriguez-Medina, J., Siebert, S., Han, D., Valderrama-Gómez, M. Á., Vahldick, H., Shulse, C. N., Cole, B. J., Juliano, C. E., Dickel, D. E., Savageau, M. A., & Brady, S. M. (2019). Molecular Mechanisms Driving Switch Behavior in Xylem Cell Differentiation. Cell Reports, 28(2), 342–351.e4. 10.1016/j.celrep.2019.06.041

Vega, S. H., Sauer, M., Orkwiszewski, J. A. J., & Poethig, R. S. (2002). The *early phase change* Gene in Maize. The Plant Cell, 14(1), 133–147. 10.1105/tpc.010406

Wang, B., Xiong, W., & Guo, Y. (2024). Dhurrin in Sorghum: Biosynthesis, Regulation, Biological Function and Challenges for Animal Production. Plants, 13(16), 2291. 10.3390/plants13162291

Wang, M., Ding, L., Gao, L., Li, Y., Shen, Q., & Guo, S. (2016). The Interactions of Aquaporins and Mineral Nutrients in Higher Plants. International Journal of Molecular Sciences, 17(8), 1229. 10.3390/ijms17081229

Wang, X., Shen, C., Meng, P., Tan, G., & Lv, L. (2021). Analysis and review of trichomes in plants. BMC Plant Biology, 21(1), 70. 10.1186/s12870-021-02840-x

Wang, Y., Zhao, Z., Liu, F., Sun, L., & Hao, F. (2020). Versatile Roles of Aquaporins in Plant Growth and Development. International Journal of Molecular Sciences, 21(24), 9485. 10.3390/ijms21249485

Wang, Y., Zhou, Q., Meng, Z., Abid, M. A., Wang, Y., Wei, Y., Guo, S., Zhang, R., & Liang, C. (2022). Multi-Dimensional Molecular Regulation of Trichome Development in Arabidopsis and Cotton. Frontiers in Plant Science, 13, 892381. 10.3389/fpls.2022.892381

Wang, Z., Yang, Z., & Li, F. (2019). Updates on molecular mechanisms in the development of branched trichome in Arabidopsis and nonbranched in cotton. Plant Biotechnology Journal, 17(9), 1706–1722. 10.1111/pbi.13167

War, A. R., Paulraj, M. G., Ahmad, T., Buhroo, A. A., Hussain, B., Ignacimuthu, S., & Sharma, H. C. (2012). Mechanisms of plant defense against insect herbivores. Plant Signaling & Behavior, 7(10), 1306–1320. 10.4161/psb.21663

Watts, S., & Kariyat, R. (2021). Morphological characterization of trichomes shows enormous variation in shape, density and dimensions across the leaves of 14 *Solanum* species. AoB PLANTS, 13(6), plab071. 10.1093/aobpla/plab071

Wu, F., Sheng, P., Tan, J., Chen, X., Lu, G., Ma, W., Heng, Y., Lin, Q., Zhu, S., Wang, J., Wang, J., Guo, X., Zhang, X., Lei, C., & Wan, J. (2015). Plasma membrane receptor-like kinase leaf panicle 2 acts downstream of the DROUGHT AND SALT TOLERANCE transcription factor to regulate drought sensitivity in rice. Journal of Experimental Botany, 66(1), 271–281. 10.1093/jxb/eru417

Wu, S., & Zhao, B. (2017). Using Clear Nail Polish to Make Arabidopsis Epidermal Impressions for Measuring the Change of Stomatal Aperture Size in Immune Response. In L. Shan & P. He (Eds.), Plant Pattern Recognition Receptors (Vol. 1578, pp. 243–248). Springer New York. 10.1007/978-1-4939-6859-6_20

Xu, P., Ali, A., Han, B., & Wu, X. (2018). Current Advances in Molecular Basis and Mechanisms Regulating Leaf Morphology in Rice. Frontiers in Plant Science, 9, 1528. 10.3389/fpls.2018.01528

Xu, Y., Qian, Z., Zhou, B., & Wu, G. (2019). Age-dependent heteroblastic development of leaf hairs in *Arabidopsis*. New Phytologist, 224(2), 741–748. 10.1111/nph.16054

Yoshikawa, T., Ozawa, S., Sentoku, N., Itoh, J.-I., Nagato, Y., & Yokoi, S. (2013). Change of shoot architecture during juvenile-to-adult phase transition in soybean. Planta, 238(1), 229–237. 10.1007/s00425-013-1895-z

Zhang, T.-Q., Chen, Y., Liu, Y., Lin, W.-H., & Wang, J.-W. (2021). Single-cell transcriptome atlas and chromatin accessibility landscape reveal differentiation trajectories in the rice root. Nature Communications, 12(1), 2053. 10.1038/s41467-021-22352-4

Zhang, T.-Q., Xu, Z.-G., Shang, G.-D., & Wang, J.-W. (2019). A Single-Cell RNA Sequencing Profiles the Developmental Landscape of Arabidopsis Root. Molecular Plant, 12(5), 648–660. 10.1016/j.molp.2019.04.004

Zhang, X., Wang, Y. P., Song, X., Zhou, L.-Z., Yu, H., Yang, L., Wang, Y. K., Wang, X. Y., Wan, X. Y., Liu, Y., Shi, Y., Yue, Z., Hou, Y., Zhang, X. S., Li, B., & Su, Y. H. (2026). A single-cell-resolution spatial transcriptomic atlas decodes wheat spike development and yield potential. Molecular Plant, 19(2), 402–424. 10.1016/j.molp.2025.12.020

Zupin, M., Sedlar, A., Kidrič, M., & Meglič, V. (2017). Drought-induced expression of aquaporin genes in leaves of two common bean cultivars differing in tolerance to drought stress. Journal of Plant Research, 130(4), 735–745. 10.1007/s10265-017-0920-x

